# *Heligmosomoides bakeri* and *Toxoplasma gondii* co-infection leads to increased mortality associated with intestinal pathology

**DOI:** 10.1101/2021.05.27.445631

**Authors:** Edina K. Szabo, Christina Bowhay, Emma Forrester, Holly Liu, Beverly Dong, Aralia Leon Coria, Shashini Perera, Beatrice Fung, Namratha Badawadagi, Camila Gaio, Kayla Bailey, Manfred Ritz, Joel Bowron, Anupama Ariyaratne, Constance A. M. Finney

## Abstract

Co-infections are a common reality but understanding how the immune system responds in this context is complex and can be unpredictable. *Heligmosomoides bakeri* (parasitic roundworm, previously *Heligmosomoides polygyrus*) and *Toxoplasma gondii* (protozoan parasite) are well studied organisms that stimulate a characteristic Th2 and Th1 response, respectively. Several studies have demonstrated reduced inflammatory cytokine responses in animals co-infected with such organisms. However, while general cytokine signatures have been examined, the impact of the different cytokine producing lymphocytes on parasite control/clearance is not fully understood. We investigated five different lymphocyte populations (NK, NKT, γδ T, CD4 ^+^ T and CD8^+^ T cells), five organs (small intestine, Peyer’s patches, mesenteric lymph nodes, spleen and liver), and 4 cytokines (IFN γ, IL-4, IL-10 and IL-13) at two different time points (days 5 and 10 post *T. gondii* infection). We found that co-infected animals had significantly higher mortality than either single infection. This was accompanied by transient and local changes in parasite loads and cytokine profiles. Despite the early changes in lymphocyte and cytokine profiles, severe intestinal pathology in co-infected mice likely contributed to early mortality due to significant damage by both parasites in the small intestine. Our work demonstrates the importance of taking a broad view during infection research, studying multiple cell types, organs/tissues and time points to link and/or uncouple immunological from pathological findings. Our results provide insights into how co-infection with parasites stimulating different arms of the immune system can lead to drastic changes in infection dynamics.

## Introduction

Co-infections are a common reality (1). However, understanding the complexities which underlie their impact on host immune responses remains difficult.

Co-infection by pathogens stimulating different arms of the immune response can result in different outcomes (2), although certain traits are thought to predict the interplay between multiple infectious agents. For example, parasite infection dynamics can be predicted by the type of immune effector mechanisms required to clear infection, as well as the ability of parasites to induce immunosuppression (3). Both these traits are defined by specific cytokine signatures. Changes in the characteristic signatures can have significant impacts on disease management. This is particularly important in the context of intestinal parasitic worm infections since these are extremely common in humans (approximately 1.8 billion infections worldwide (4)), livestock (5) and wildlife (6). The parasites negatively impact vaccine and treatment efficacy (7,8), which can be restored with effective anthelminthic treatment that clear the worms (9).

In real-world infections, the intestinal mucosa is continually subjected to a multitude of pathogens and commensals. The interplay between the different organisms impacts the ability of the immune system to maintain a strong barrier and contain these populations. The gut-associated lymphoid tissue (GALT), including the mesenteric lymph nodes (MLN), Peyer’s Patches (PP) and the small intestine (SI), represents the largest lymphatic mass in the body. The ability of lymphocytes to traffic to and interact at these sites is critical to host defence (10). Here, we studied the impact of two parasites that stimulate opposing immune responses in the small intestine and the GALT.

*Heligmosomoides bakeri* (Hb), formerly *Heligmosomoides polygyrus* (11), is a parasitic nematode (roundworm) which matures within the intestinal tissue and causes a chronic infection whereby adult worms reproduce and lay eggs in the intestinal lumen (12). The immune response to this parasite has been well characterised (12). To clear worms, the host stimulates a potent Th2 immune response leading to the formation of a Th2 granuloma that allows the trapping/killing of larvae (13). Antibodies and intestinal physiological responses also promote worm killing/expulsion. In conjunction, a Treg response is mounted, thought to limit any potentially immunopathological Th1 response (14). However, in susceptible mice (C57Bl/6 mice, used here), the granulomas and relatively weak Th2 response are not strong enough to clear the worms (15); the infection remains chronic.

*Toxoplasma gondii* (Tg) is an intracellular parasite that first infects intestinal cells, and multiple other cell types as it spreads systemically (16). The host response to Tg is a strong inflammatory response involving IFNγ production by lymphocytes. NK cells are essential for the early control of Tg (17–19), as is the induction of CD8^+^ T cells (20). Similarly, Tg infection studies demonstrate that γδ T cell depleted mice are less resistant to Tg than their wildtype counterparts, due to the early IFN γ production of γδ T cells during infection (21,22). The role of NKT cells during Tg infection is not clear. While they play a large part in the induction and initiation of inflammatory responses, this can lead to overproduction of IFN γ, which can be detrimental for the host (23). Others have found that NKT cells may even be directly involved in the suppression of protective immunity against Tg (24). However, most studies published on helminth-Tg co-infections focus on IFN γ production by conventional T cells (CD4 ^+^ and CD8 ^+^ T cell) (25–28), with much less known about the role of other lymphocyte populations (e.g. γδ T and NKT cells). Finally, as well as a strong IFNγ response, Tg stimulates high levels of IL-10 to limit immunopathology(29).

Several studies have demonstrated reduced inflammatory cytokine responses in animals co-infected with an intestinal nematode and a microparasite such as Tg (26,28,30–32). While general cytokine signatures have been examined, the impact on the different cytokine producing lymphocytes on parasite control/clearance is not fully understood. To shed more light on the lymphocyte response in the context of HbTg co-infection, we investigated the changes in Th1 and Th2 cytokines at the systemic level, as well as IFN γ-producing lymphocytes (NK, NKT, CD4 ^+^ T, CD8 ^+^ T and γδ T cells) at day 5 and day 10 post-Tg infection in different organs. Few studies have focused on the SI, which is Hb’s niche and is Tg’s first contact with the host, or on the PP which are intricately linked to the SI. This is important since lymphocytes generally display distinct phenotypes and functions in different organs(33).

Here, we found that co-infected animals had significantly higher mortality than either single infection. This was accompanied by changes in cytokine profiles and parasite loads which were dependent on the cell type (CD4^+^ T, CD8^+^ T, NK, NKT and γδ T cells), the organ (SI, PP, MLN, SPL and liver), the time-point (day 5 and 10 post-infection) and the cytokine (IFNγ, IL-4, IL-10 and IL-13) studied. These changes do not fully account for the mortality differences between the groups. However, our pathology results indicate that HbTg infected animals suffered from significant damage by both parasites in the SI, which was likely a significant contributor to increased mortality.

## Methods

### Animals, parasites and infection protocols

C57BL/6 female mice, aged between 8-10 weeks old, were bred in house at the University of Calgary. Breeding pairs were originally purchased from Charles River Laboratories (Senneville, Quebec). All mice were housed under specific pathogen-free conditions. The University of Calgary’s Animal Care Committee approved all experimental animal procedures.

The Me49 strain of *Toxoplasma gondii* (Tg) (maintained in house, original stock was a gift from Dr. Georgia Perona Wright, University of British Columbia, Canada) was maintained in male C57BL/6 mice by bimonthly passage into new animals. Cysts were obtained from the brains of these animals a month after infection. Brains of infected mice were homogenised in PBS to count tissue cysts. 20 cysts were orally administered per mouse (female C57BL/6). Animals were euthanized either 5 or 10 days post infection.

Female C57BL/6 mice were orally infected with 200 *Heligmosomoides polygyrus bakeri* (Hb) infective larvae (maintained in house, original stock was a gift from Dr. Allen Shostak, University of Alberta, Canada). Animals were euthanized either 12 or 17 days post infection. Larvae were obtained from fecal cultures after approximately 8-9 days of incubation at room temperature.

For co-infected animals, mice were first orally infected with Hb for 7 days. At day 7 post-infection, animals were orally infected with 20 cysts of the Me49 strain of Tg (all other experimental animals were gavaged with PBS as a control). Mice were euthanized 5 or 10 days post Tg infection (equivalent to 12 or 17 days post Hb infection).

For worm counts, small intestines of Hb and HbTg infected mice were harvested and opened longitudinally. The number of adult worms present in the intestinal lumen along the length of the small intestine was counted under a dissection microscope.

### Histopathology

Small intestines, spleens and livers were isolated and fixed in 4% paraformaldehyde. Paraffin embedded sections were obtained for each tissue and stained with hematoxylin and eosin (H&E) (for eosinophil/macrophage identification in the granulomas) and/or hematoxylin and alcian blue (for goblet cell counts in the small intestine). Sections were cut and stained with H&E by the University of Calgary’s Veterinary Diagnostic Services Unit. Stained tissue sections were analysed visually using the Leica DMRB light microscope and/or the OLYMPUS SZX10 microscope according to a scoring system developed by the Finney group (S1 and S2 Fig, S1 and S2 Tables). The intestinal inflammation score was adapted from (34). Images were captured using the QCapture and the CellSens software and processed using the ImageJ software. For eosinophil and macrophage identification, one photograph of each granuloma was taken at x400 magnification using an Olympus BX 34 microscope and the Olympus CellSens software. Cells were identified by their morphology and staining patterns.

For the small intestinal, spleen and liver scoring systems, slides were blindly evaluated by two independent researchers at x40 magnification. Three scoring parameters were developed for the spleen: follicle shape, marginal zone thickness and size of the germinal center (light zone). For the liver, we used the presence/absence of necrosis, infiltrate size, proportion of infiltrates, perivascular infiltrate characteristics. The parameters within each scheme were given an equal weighting and were combined to give a total, gross pathological score for each slide. A higher score is representative of a greater amount of observed tissue damage. The pathology scale for the small intestine was adapted from (35).

For paneth cell counts, the total number of cells per 20 VCU (villus/crypt unit) was calculated in the proximal and distal SI for each animal. For goblet cell counts, the average number of goblet cells from five consecutive villi was calculated for each animal in the proximal and distal SI. The villi perimeters were calculated using ImageJ.

### Serum ELISAs

Blood samples were collected using a terminal cardiac bleed. Blood was left to clot for 30 minutes and then centrifuged twice at 11, 000 g at 4°C for 10 minutes.

IFNγ cytokine production was measured in the serum of Tg and HbTg infected, and naive mice with the Mouse DuoSet ELISA development system (R&D Systems), according to the manufacturers’ instructions.

### RNA/DNA Extraction and Quantitative PCR

RNA and gDNA were extracted from snap frozen SPL, LIV, MLN, PP, and 4 equal sections of the SI (sections 1-4, where 1 is the most proximal section and 4 the most distal from the stomach) by crushing the tissue using a pestle and mortar on dry ice and using Trizol (Ambion, Life Technologies) following manufacturer guidelines.

To measure cytokine levels in the RNA samples, cDNA was prepared using Perfecta DNase I (Quanta), and qScript cDNA Supermix (Quanta) or iScript Reverse Transcription Supermix for RT-qPCR (Bio-Rad).

Primers for quantitative PCR were obtained from Integrated DNA Technologies (San Diego). For IFNγ TCAAGTGGCATAGATGTGGAAGAA forward, and TGGCTCTGCAGGATTTTCATG reverse, IL-10 GTCATCGATTTCTCCCCTGTG forward, and ATGGGCCTTGTAGACACCTTG reverse, IL-4 CGAAGAACACCACAGAGAGTGAGCT forward, and GACTCATTCATGGTGCAGCTTATC reverse, IL-13 GATCTGTGTCTCTCCCTCTGA forward, and GTCCACACTCCATACCATGC reverse, ZO-1 CGCCAAATGCGGTTGATC forward and TTTACACCTTGCTTAGAGTCAGGGTT reverse, Occludin CCAGGCAGCGTGTTCCT forward and TTCTAAATAACAGTCACCTGAGGGC reverse, IL-22 ACTTCCAGCAGCCATACATC forward and CACTGATCCTTAGCACTGACTC reverse and β-actin GACTCATCGACTCCTGCTTG forward, and GATTACTGCTCTGGCTCCTAG reverse primers were used. To confirm primer product size, all products were run on a gel. Samples were run on a Bio-Rad CFX96 real time system C1000 touch thermocycler. Relative quantification of the cytokine genes of interest was measured with the delta-delta cycle threshold quantification method (36), with β-actin for normalization. Data are expressed as a fold change relative to uninfected samples.

To quantify the *T. gondii* parasite burdens in the gDNA samples, we used a standard curve of *T. gondii* tachyzoites. *T. gondii* B1 was the target gene for the quantification of parasite in all tissues or organs (37). B1 primers (forward: TCCCCTCTGCTGGCGAAA, reverse: AGCGTTCGTGGTCAACTATCGATT) were prepared by IDT (San Diego).

### Flow Cytometry

Cell suspensions were obtained from the MLN, and LIV. For the MLN, cells suspensions were obtained through mechanical disruption of whole tissues using a pestle. Perfused liver lobes were homogenised as per MLN. Perfusion was carried out through the vena cava after mice were anaesthetised, using a peristaltic pump (Gilson Minipuls3) to circulate prewarmed PBS until a change of colour was observed in the liver. Lymphocytes were isolated from the cell suspension using a 37 % and 70 % percoll gradient (GE Healthcare Bio-Sciences AB, Sweden) diluted with HBSS (with or without phenol red, Lonza, Switzerland), and resuspended in RPMI.

For *ex vivo* restimulation, cells were incubated with 50ng/mL phorbol myristate acetate and 1 μg/mL ionomycin for 6 h at 37 °C with 5% CO _2_ in the presence of BD GolgiStop. Single cell suspensions for all tissue types were stained and analysed according to (38). Cells were stained for: viability (APC-H7 or AF700, BD Biosciences), surface markers: CD45 (BV510), CD3 (BV605), CD4 (PerCP-Cy5.5), CD8 (BUV395), NK1.1 (BV421), γδ TCR (BV711) and intracellular markers: IFNγ (APC), and granzyme B (FITC). All antibodies were purchased from BD Biosciences apart from CD4 (PerCP-Cy5.5, BioLegend), Granzyme B (FITC, eBioscience). Cells were blocked with rat α-mouse CD16/32 (Biolegend). Cells were run on an LSRFortessa X-20 flow cytometer and data analysed using FlowJo software. For analysis, doublets and dead cells were removed from the analysis, and NK, NKT, γδ T, CD4 T and CD8 T IFNγ and GZMB producing cells were quantified (S3 Fig).

### Statistical Analysis

Graphpad Prism software (La Jolla, CA, USA) was used for all statistical analysis. To compare multiple groups (three or more), we performed a normality test (D’Agostino & Pearson test, unless N too small, in which case Anderson-Darling or Shapiro-Wilk). ANOVA or Kruskal-Wallis tests were performed on parametric/non-parametric pooled data, and when significant, Sidak’s/Dunn’s Multiple comparisons were performed on Hb vs. HbTg and Tg vs. HbTg. For survival analysis, we used a Mantel-Cox test. Data are presented as median and individual data points, unless otherwise specified.

## Results

### Increased mortality of HbTg infected mice correlates with increased intestinal worm burden and MLN Tg burden

200 *H. bakeri* larvae (L3) were orally administered to female C57Bl/6 mice 7 days prior to infection with 20 Me49 *T. gondii* tissue cysts (Fig 1A). By day 16, over 60% (8/12) Tg-infected mice survived infection in contrast to fewer than 20% (2/12) HbTg-infected animals (Fig 1B).

**Fig 1:**
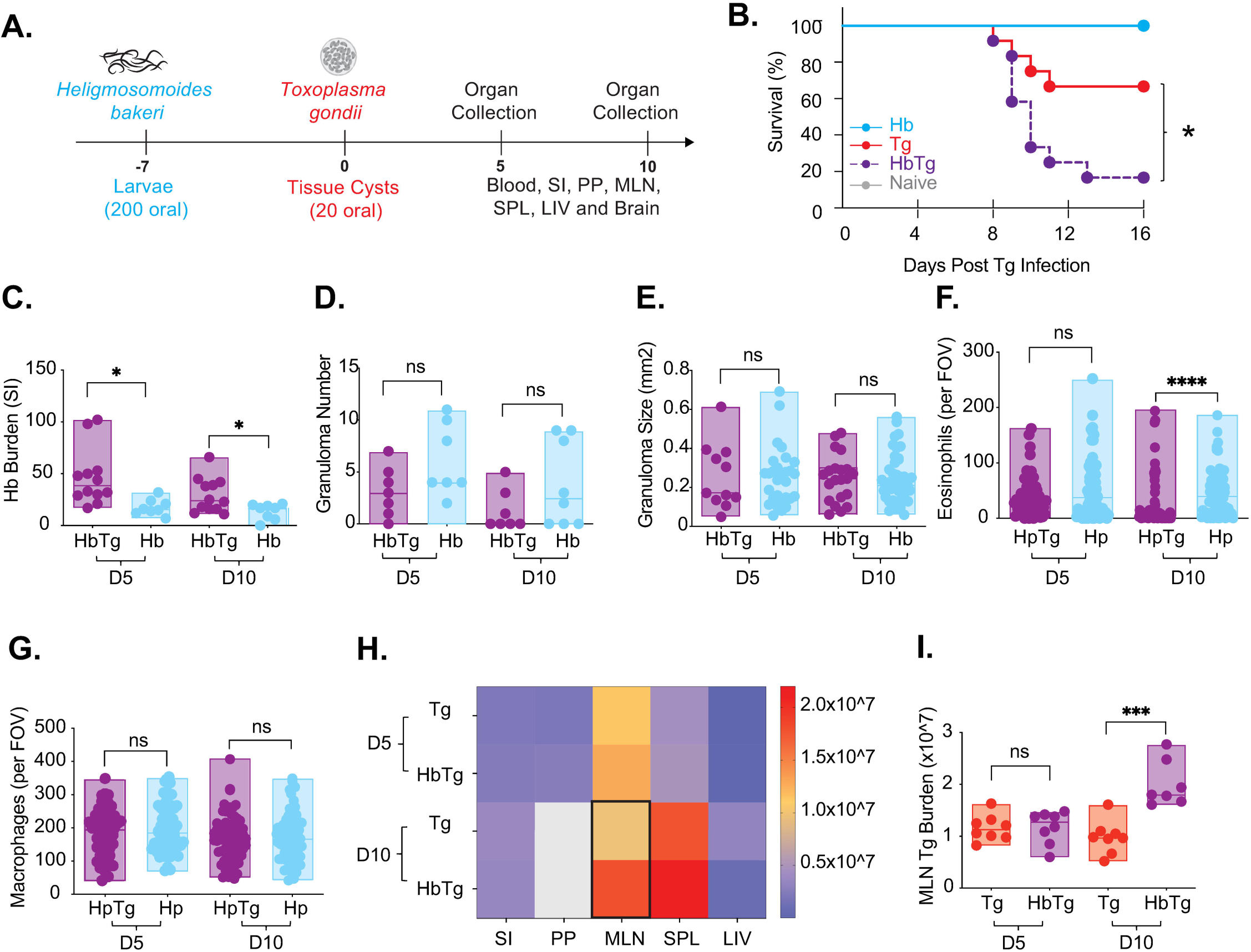
*H. bakeri* co-infection leads to high mortality and increased parasite loads in mice co-infected with *T. gondii*. (A) 200 Hb larvae were given orally to mice 7 days prior to infection with 20 Me49 Tg tissue cysts. Animals were euthanised 5 and 10 days post Tg infection (12 and 17 days post Hb infection). (B) Survival was measured for 16 days post Tg infection. N = 4 mice (Tg and HbTg-infected groups) and 2-4 mice (naïve/Hb-infected groups) per experiment, 3 independent experiments. Mantel-Cox test performed for pooled data, *p=0.02 (Tg vs. HbTg). Adult worms (C) and granulomas (D) were counted in the small intestine of Hb and HbTg infected mice at 12 and 17 days post Hb infection. (E) Granuloma size was calculated for the granulomas of Hb and HbTg infected mice at 12 and 17 days post Hb infection. The perimeter of each granuloma was measured and the surface area calculated (a minimum of 11 granulomas measured per group). Eosinophil (F) and macrophage (G) counts within the centre of the granulomas. Formalin fixed, paraffin-embedded 6 μm sections were obtained from whole small intestine swiss rolls. H & E slides were used to identify eosinophils and macrophages at x400 magnification. Cells were counted for one field of view (FOV) per granuloma (a minimum of 55 granulomas per group). (H & I) Tg loads were measured by quantitative PCR in the small intestine (SI), Peyer’s patches (PP), mesenteric lymph nodes (MLN), spleen (SPL) and liver (LIV) at 5 and 10 days post-Tg-infection. Grey boxes depict values that are out of range (>2.0*10^7). Black outline depicts significant differences between Tg and HbTg groups. For (C-I), N= 2-6 mice per group per experiment, 2 independent experiments. All data were tested for normality. Unpaired T-tests/Mann Whitney tests were performed on parametric/non-parametric pooled data on either Hb vs. HbTg or Tg vs. HbTg; n.s. = non significant, *= p<0.05, ***= p<0.001 and ****= p<0.0001.

To determine whether decreased survival was associated with increased parasite loads, we measured Hb and Tg levels. Tg and Hb both infect their host via the small intestine. Hb has an entirely enteric lifecycle. Larval stages develop within the intestinal tissue resulting in the formation of granulomas (accumulation of immune cells around the worm), which are visible for weeks even after the worms have been cleared. Increased granuloma numbers and size are associated with resistance (15). Adult stages (found at days 12 and 17 post-infection) reside in the intestinal lumen (39). At both time points, we found an increase in adult worms in the lumen of HbTg compared to Hb infected mice (Fig 1C) but no difference in granuloma number (Fig 1D) or size (Fig 1E). We found very few granulomas (no more than two) containing developing worms at days 12 and 17 post Hb infection. Numbers were not different between HbTg and Hb infected animals (data not shown), suggesting that differences in worm infection kinetics were not responsible for the differences in worm burden observed between the two groups. Granulomas did not differ in their composition 5 days post infection, with Tg and HbTg infected animals having similar numbers of eosinophils and macrophages (Fig 1E & 1F). By 10 days post-infection, the number of eosinophils was reduced in the HpTg group (Fig 1F).

Unlike Hb, Tg infects its host through the small intestine, and quickly disseminates throughout the body. This can occur as early as 3 days post-infection (40–42). We measured Tg levels in the SI, PP, MLN, SPL and LIV at both 5 and 10 days post Tg infection. Tg was detected in all organs, at both time points in Tg and HbTg infected animals. At day 5 post Tg infection, the organ with the highest level of Tg were the MLN (Fig 1F). No difference in Tg burden between Tg and HbTg infected animals was observed at this time point in any of the organs (Fig 1F & 1G). However, 10 days post Tg infection, levels in all three lymphoid organs (PP, MLN and SPL) were highest (Fig 1F) and Tg levels were increased in the MLN of HbTg compared to Tg infected mice (Fig 1G).

Commonly Me49 infection culminates in brain tissue being invaded by approximately 3 weeks post-infection. No tissue cysts were found in any of the infected mice at d5 or day 10 post-infection, or at the time of death (up to day 16 post-infection).

### Cytokine profiles differ between Tg and HbTg infected mice in the MLN

We measured cytokine levels in the MLN to assess whether they were associated with the increased Tg burden measured on day 10 post infection in HbTg compared to Tg infected animals.

Th1 responses, specifically IFN γ produced by NK and CD8 ^+^ T cells, are crucial for protection against Tg (43–49). A lack of early IFNγ leads to increased Tg replication (50,51). At 5 days post Tg infection, HbTg animals had only approximately a third of the *Ifnγ* expression measured in the Tg-infected group (Fig 2A). By 10 days post Tg infection, levels in Tg animals had decreased and the difference between the two groups was no longer apparent (Fig 2A). We also measured *Ifnγ* expression levels in the organs where we measured Tg burden (Fig 2B). Since Tg is mostly associated with pathology in the ileum (52,53), but has also been found to replicate along the entire SI (both proximal and distal (42)), and Hb adult worms are mostly found in the proximal part of the SI (in close proximity to the stomach (54)), we analysed gene expression in the SI, divided into four equal parts (SI1-4, from proximal to distal). We also measured systemic IFN γ protein levels in the serum (S4 Fig). As expected, levels of *Ifnγ* expression in naïve and Hb infected animals were low, but elevated in the Tg infected animals (Fig 2B). The MLN were the organ with the highest fold increase (approximately 700, 5 days post Tg infection, Fig 2B) and the only site with a significant change in expression between Tg and HbTg infected animals. This difference did not translate to differences in serum protein levels (S4 Fig).

**Fig 2:**
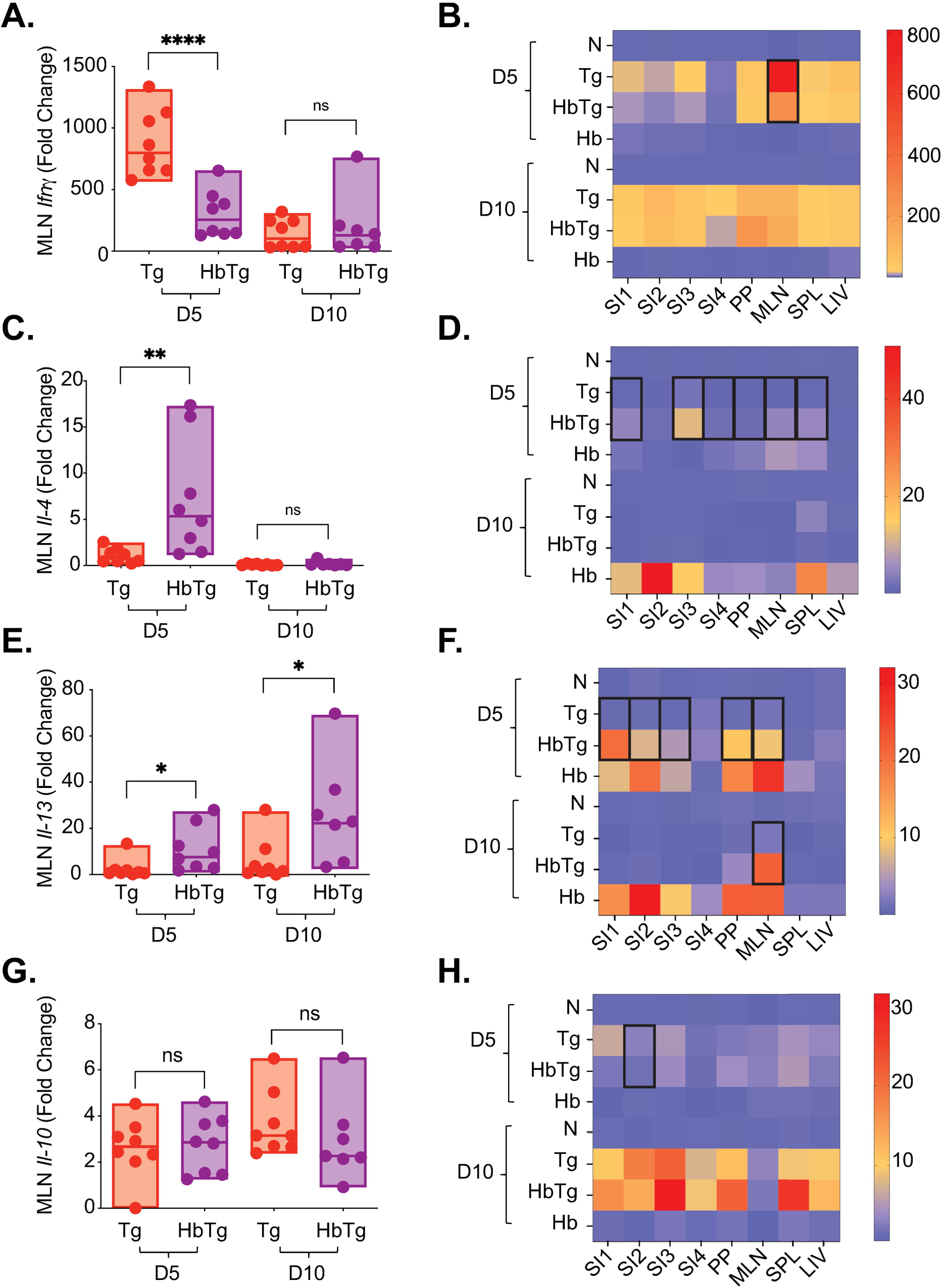
The MLN gene expression profile in co-infected mice differs from that in *T. gondii*-infected mice. 200 Hb larvae were given orally to mice 7 days prior to infection with 20 Tg tissue cysts. Gene expression was measured by quantitative RT-PCR at 5 and 10 days post-Tg-infection for *Ifnγ* (A), *Il-4* (C), *Il-13* (D) and *Il-10* (E) in the mesenteric lymph nodes (MLN). Heat maps represent median levels of *Ifnγ* (B), *Il-4* (D), *Il-13* (E) and *Il-10* (F) in the small intestine (SI, regions 1-4 from duodenum to ileum), Peyer’s patches (PP), MLN, spleen (SPL) and liver (LIV) at 5 and 10 days post-Tg-infection. Black outline depicts significant differences between Tg and HbTg. N= 2-4 mice per group per experiment, a minimum of 2 independent experiments. Data was tested for normality. ANOVA or Kruskal-Wallis tests were performed on parametric/non-parametric pooled data for N/Tg/HbTg/Hb, and when significant, Sidak’s/Dunn’s Multiple comparisons were performed on Tg vs. HbTg; n.s. = non significant, * = *p*<0.05, ** = *p*<0.01 and ****= *p*<0.0001.

The production of Th2 cytokines can lead to reduced inflammatory responses (55,56), with IL-4 production directly inhibiting IFN γ production (57). The Th2 cytokines IL-4, IL-13 and IL-10 are involved in worm expulsion (15,58–60). In the absence of IL-4 and IL-13, worm expulsion is compromised (60) and high levels of IL-10 have been associated with resistant phenotypes (15). IL-10 expression is also required in the SI to prevent necrosis and mortality of Tg infected animals (61). We measured the gene expression profiles of *Il-4*, *Il-13* and *Il-10* (Fig 2C - H). Unsurprisingly, we found elevated *ll-4* and *Il-13* expression in the HbTg compared to Tg infected animals at day 5 post Tg infection in the MLN (Fig 2C & 2E). This increase was still apparent for *Il-13* 10 days post Tg infection (Fig 2E). *Il-10* expression levels did not differ between the two groups at either time point (Fig 2G).

As expected, levels of *Il-4* and *Il-13* were increased in all organs tested in Hb infected animals, and to a much higher extent than in Tg infected animals (Fig 2D & 2F). In general, the *Il-4* and *Il-13* gene expression profiles of co-infected animals were more similar to Hb animals at day 5 post Tg infection. For example, in the SI, *Il-4* and *Il-13* production were increased in HbTg compared to Tg infected animals in at least 3 of the 4 sections (Fig 2D & 2F). By 10 days post infection, this was no longer the case. Despite the reduced Hb loads in the HbTg vs. Hb animals, we observed no differences in *Il-13* cytokine gene expression between the HbTg and Hb animals, and only one difference in *Il-4* in I3 at day 5 post Tg infection.

### In the MLN, decreased IFN**γ** production is observed across five cell types but is not associated with decreased granzyme B production

The MLN are the draining lymph nodes to the SI. Different lymphocytes produce IFN γ within this immune organ, including NK and CD8 ^+^ T cells, traditionally associated with IFN γ production during Tg infection (62–65), as well as NKT, γδ and CD4 ^+^ T cell more recently implicated in the inflammatory response to Tg infection (66–68). To determine whether a particular cell type was associated with the decreased IFN γ production observed in HbTg in the MLN at 5 days post-Tg infection, we studied cell kinetics and IFN γ production in these five cell types. IFN γ-producing cells are mainly observed in the Tg-infected group (Fig 3A). γδ T and NK cells have the highest proportion of IFN γ producers with a median of approximately 12% each. Across all cell types, the percentage IFN γ-producing cells is similar between HbTg and Hb-infected animals (Fig 3A), and decreased compared to Tg-infected animals (Fig 3A-H). By 10 days post-Tg infection, the proportion all IFN γ producing lymphocytes studied in the MLN of HbTg infected animals were no longer decreased compared to those in Tg-infected mice (data not shown).

**Fig 3:**
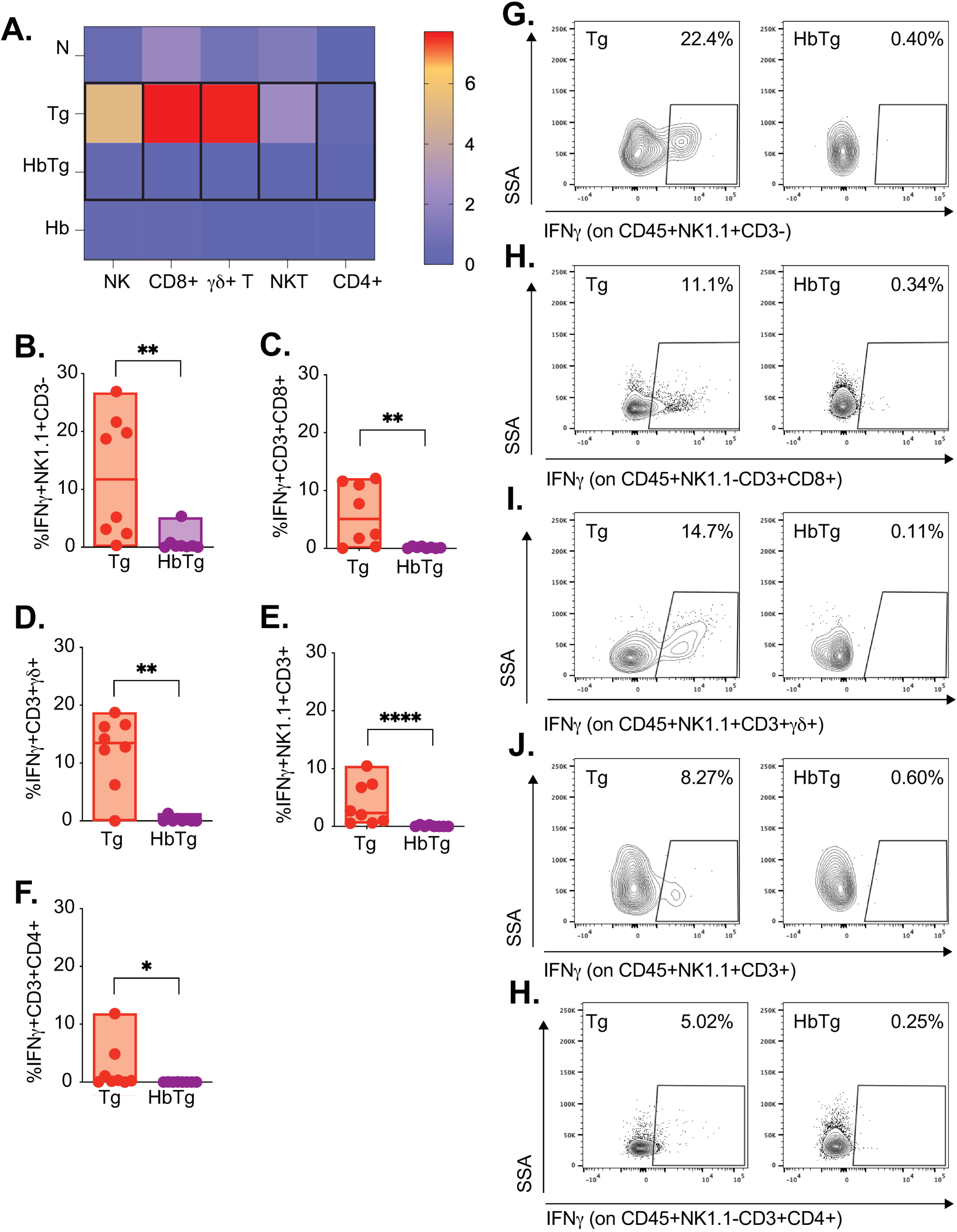
Co-infected mice have reduced levels of MLN IFNγ-producing NK, CD8^+^ T, γδ T, NKT and CD4^+^ T cells. 200 Hb larvae were given orally to mice 7 days prior to infection with 20 Tg tissue cysts. (A) Heat map represents median levels of IFNγ producing cells in the MLN at 5 days post-Tg-infection. Black outline depicts significant differences between Tg and HbTg. The cell percentage of IFNγ-producing (B) NK (IFNγ+NK1.1^+^CD3-), (C) CD8 T (IFNγ+CD3^+^CD8^+^), (D) NKT (IFNγ+NK1.1^+^CD3^+^), (E) γδ T (IFNγ+ CD3^+^γδ+) and (F) CD4 T cells (IFNγ+CD3^+^CD4^+^) in Tg and HbTg animals. (G-H) Flow plots depicting the IFNγ+ cells in Tg and HbTg animals. N= 2-4 mice per group per experiment, 2 independent experiments. Data was tested for normality. ANOVA or Kruskal-Wallis tests were performed on parametric/non-parametric pooled data including N/Tg/HbTg/Hb groups, and when significant, Sidak’s/Dunn’s Multiple comparisons were performed on Hb vs. HbTg and Tg vs. HbTg; n.s. = non significant, * = *p*<0.05, ** = *p*<0.01 and ****= *p*<0.0001.

Interestingly, Granzyme B production did not mirror IFN γ production (GZMB, Fig 4). GZMB, a serine protease that activates apoptosis, has traditionally been associated with NK and CD8 ^+^ T cell killing mechanisms, explaining its increase during Tg infection (69). However, the extent of its role during Tg infection remains controversial. Some studies have shown that Tg can actively inhibit GZMB in infected cells (70) implying that GZMB production negatively impacts Tg infection. Others have found GZMB to have a limited role in host protection (71). GZMB is also upregulated during nematode infections, although its precise role (harmful/beneficial to the host) also remains controversial (72). We observed no change in the proportion of GZMB-producing lymphocytes between the HbTg and Tg infected groups in any of the cell types, with NK cells having the largest proportion of GZMB producers (Fig 4A-F). Interestingly, we did observe an increase in GZMB production (GZMB MFI) in the NK, CD4 ^+^ T and CD8 ^+^ T cells of Tg and HpTg infected animals compared to naïve animals; differences between Tg and HpTg groups were only apparent for the CD4^+^ and CD8^+^ T lymphocytes (Fig 4G-L).

**Fig 4:**
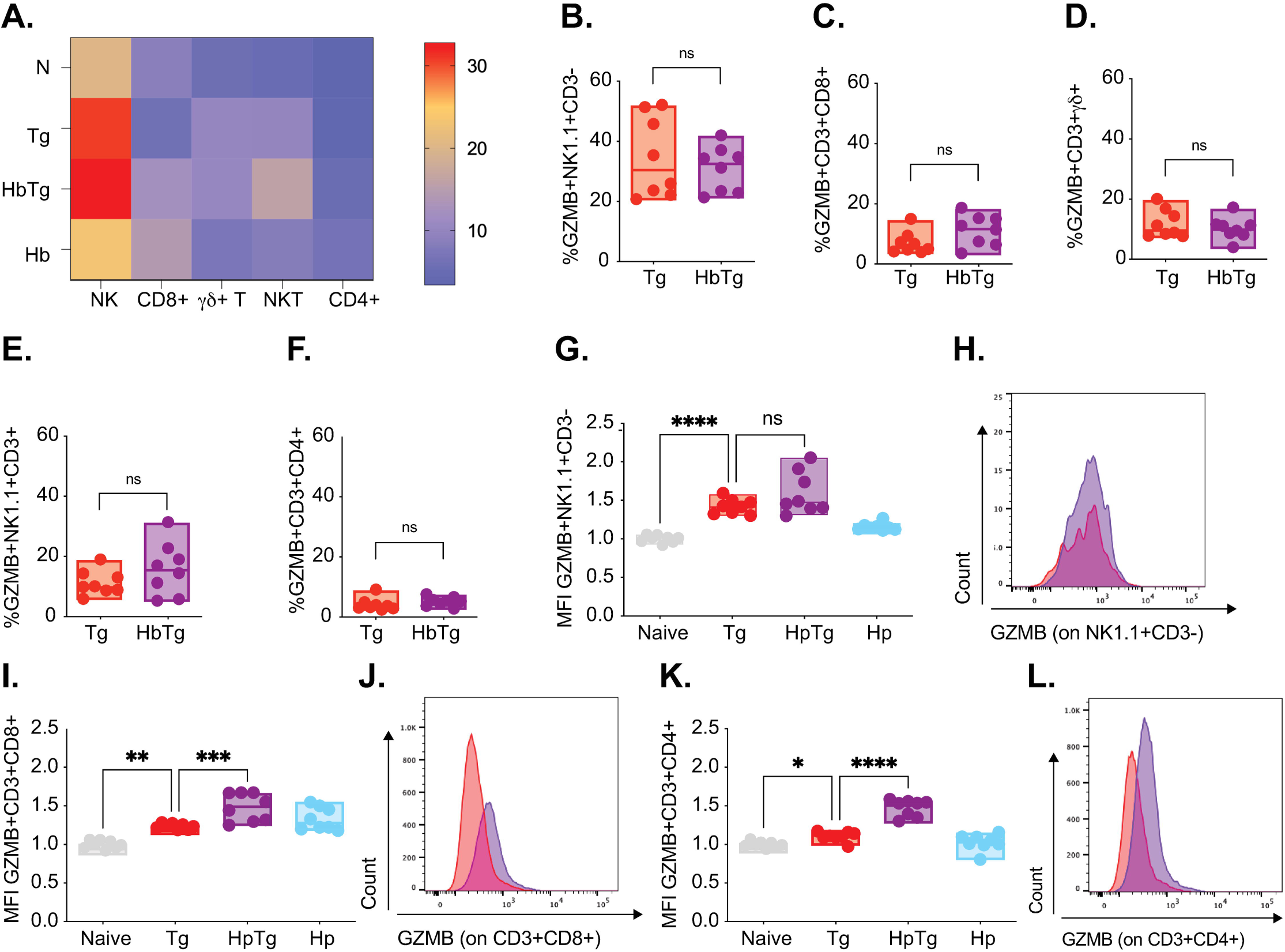
Co-infected mice have increased CD8^+^ T and CD4^+^ T cell GZMB production in the MLN. 200 Hb larvae were given orally to mice 7 days prior to infection with 20 Tg tissue cysts. (A) Heat map represents median levels of GZMB producing cells in the MLN at 5 days post-Tg-infection. The cell percentage of GZMB -producing (B) NK (GZMB+NK1.1^+^CD3-), (C) CD8 T (GZMB+CD3^+^CD8^+^), (D) NKT (GZMB+NK1.1^+^CD3^+^), (E) γδ T (GZMB+ CD3^+^γδ+) and (F) CD4 T cells (GZMB+CD3^+^CD4^+^) in Tg and HbTg animals. GZMB MFI levels on (G) NK (NK1.1^+^CD3-), (I) CD8 T (CD3^+^CD8^+^) and (K) CD4 T (CD3^+^CD4^+^) cells relative to naïve animals. Flow histograms depicting t GZMB MFI levels on (H) NK (NK1.1^+^CD3-), (J) CD8 T (CD3^+^CD8^+^) and (L) CD4 T (CD3^+^CD4^+^) in Tg and HbTg animals. N= 2-4 mice per group per experiment, 2 independent experiments. Data were tested for normality. ANOVA or Kruskal-Wallis tests were performed on parametric/non-parametric pooled data including N/Tg/HbTg/Hb groups, and when significant, Sidak’s/Dunn’s Multiple comparisons were performed on Naive vs. Tg and Tg vs. HbTg; n.s. = non significant, * = *p*<0.05, ** = *p*<0.01, *** = *p*<0.001 and ****= *p*<0.0001.

### In the PP, changes in IFN**γ** and granzyme B production are restricted to the CD8**^+^** and CD4**^+^** T cells

The PP are loosely organised lymphoid follicles found along the SI that exhibit differential responses after Hb infection (73). Tg is also known to infect and replicate within them (74). Unlike in the MLN, we found no difference between the levels of *Ifnγ* transcripts in the Tg and HbTg groups (Fig 2B). However, different levels of production by the different lymphocyte populations could account for this. We therefore studied the percentage of IFN γ-producing lymphocytes within the PP (NK, CD8 ^+^ T, NKT, γδ T and CD4 ^+^ T). Only the IFN γ-producing CD8^+^ T cells were decreased in the HbTg group compared to the Tg group at 5 days post Tg infection (Fig 5A-C). Unlike in the MLN, GZMB-producing CD8 ^+^ T cells were increased in the HbTg compared to the Tg infected groups (Fig 5D-F). And, we also observed an increase in GZMB production (GZMB MFI) in the NK and γδ (but not the αβ CD4^+^ and CD8^+^) T cells of Tg and HpTg infected animals compared to naïve animals; differences between Tg and HpTg groups were only apparent for the CD4 ^+^ T lymphocytes. By d10 post-Tg infection, no differences were observed in the PP between the HbTg and Tg groups in any of the parameters studied (data not shown). Differences between the HbTg and Tg groups were not found in any of the IFN γ-producing or the GZMB-producing lymphocytes in the spleen and/or liver at either time point (data not shown).

**Fig 5:**
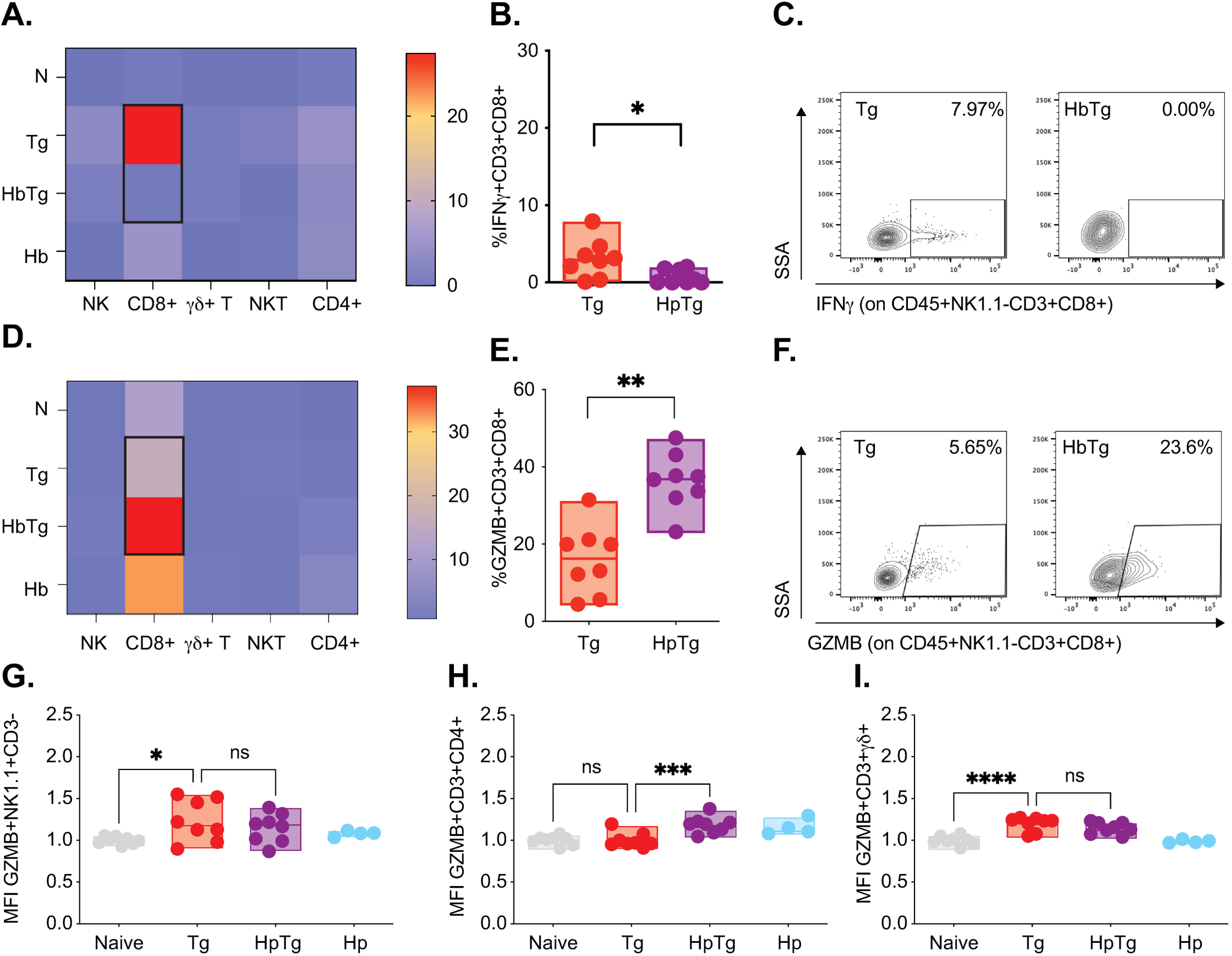
Co-infected mice have reduced levels of IFNγ-producing CD8^+^ T and increased levels of MLN GZMB-producing CD8^+^ T in the PP. 200 Hb larvae were given orally to mice 7 days prior to infection with 20 Tg tissue cysts. (A) Heat map represents median levels of IFNγ producing cells in the MLN at 5 days post-Tg-infection. (B) The cell percentage of IFNγ producing CD8 T (IFNγ+CD3^+^CD8^+^) in Tg and HbTg animals. (C) Flow plots depicting the IFNγ+ cells in Tg and HbTg animals. (D) Heat map represents median levels of GZMB producing cells in the MLN at 5 days post-Tg-infection. (E) The cell percentage of GZMB producing CD8 T (GZMB+CD3^+^CD8^+^) in Tg and HbTg animals. (F) Flow plots depicting the GZMB^+^ cells in Tg and HbTg animals. GZMB MFI levels on (G) NK (NK1.1^+^CD3-), (H) γδ T (CD3^+^γδ^+^) and (I) CD4 T (CD3^+^CD4^+^) cells relative to naïve animals. N= 2-4 mice per group per experiment, 2 independent experiments. Data were tested for normality. ANOVA or Kruskal-Wallis tests were performed on parametric/non-parametric pooled data including N/Tg/HbTg/Hb groups, and when significant, Sidak’s/Dunn’s Multiple comparisons were performed on Naive vs. Tg and Tg vs. HbTg; n.s. = non significant, * =*p*<0.05, ** = *p*<0.01, *** = *p*<0.001 and ****= *p*<0.0001.

Cell numbers, rather than subset percentages, can provide useful insights when studying organs that change size during infection. Compared to the naïve group, MLN cell numbers were increased approximately 4-5 times in all infected groups (data not shown). We did not study the cell numbers for the PP, as they are not a discreet organ and the number of cells varies per experiment. The numbers of NK, NKT, γδ T and CD4 T cells, but not CD8 T cells, were increased in the Tg infected compared to co-infected group (Fig 5A-I). This translated into a decrease in the total number of IFN γ^+^ CD8^+^ T, NKT, γδ+ T and CD4 ^+^ T cells (Fig 5F-J) and an increase in the number of GZMB^+^ NK, NKT and CD4^+^ T cells (Fig 5K-O).

### Intestinal pathology differs between Tg and HbTg infected animals

Increased Tg burden in the MLN of HbTg infected mice is associated with decreased levels of IFNγ by all IFN γ-producing lymphocytes. However, these transient, localised changes alone are unlikely to account for the difference in mortality between the HpTg and Tg infected animals. We therefore investigated whether the increased mortality and altered cytokine levels observed in co-infected animals correlated with increased pathology. We focused first on the SI, and particularly the proximal region, since this is a preferred niche for both Hb and Tg parasites.

Whole small intestines were formalin fixed, paraffin embedded and stained with hematoxylin and eosin to investigate intestinal pathology. Gene expression of tight junction proteins *Zo1* and *Occludin*, and the key regulator of inflammation *Il-22* were measured in the SI (SI2 region). Epithelial barrier dysfunction is characterised by decreased levels of *Zo1* and *Occludin*, which has been observed during Tg infection both at the gene expression (75) and protein levels (76). Transgenic animals lacking *Il-22* infected with Tg have higher mortalities than their wildtype counterparts (77). As expected, inflammation score and *Il-22* levels increased with Tg infection while *Zo1* and *Occludin* levels decreased. However, despite differences in mortality (Fig 1B), we saw no difference between Tg and HbTg groups in these parameters at day 5 (Fig 6A-E, S1 Table) or 10 (data not shown) post Tg infection. Paneth cells have recently been shown to play an important role in limiting immunopathology and microbiota dysbiosis driven by Tg infection (78,79). However, again, we found no difference between Tg and HbTg animals in paneth cell numbers at 5 (Fig 6F & 6G) or 10 (data not shown) days post Tg infection, or their state of degranulation (data not shown). The only differences we observed between Tg and HbTg infected animals in the SI were the intestinal villi size, the number of goblet cells, the weight of the intestinal tissue and the presence of granulomas. We, like others (80), observe that villus length decreases in the proximal SI during Tg infection (Fig 6H). Goblet cells serve an important physical barrier against pathogens through the maintenance and production of mucus layer in the small intestine by the production of mucins (81). Their increase in number coupled to granuloma formation, as well as the resulting increase in intestinal weights, are all associated with Hb infection (82). These changes are also observed in the HbTg, but not the Tg animals at 5 days post Tg infection (Fig 6J-L). By 10 days, the only remaining difference between Tg and HbTg infected animals is the presence of granulomas.

**Fig 6:**
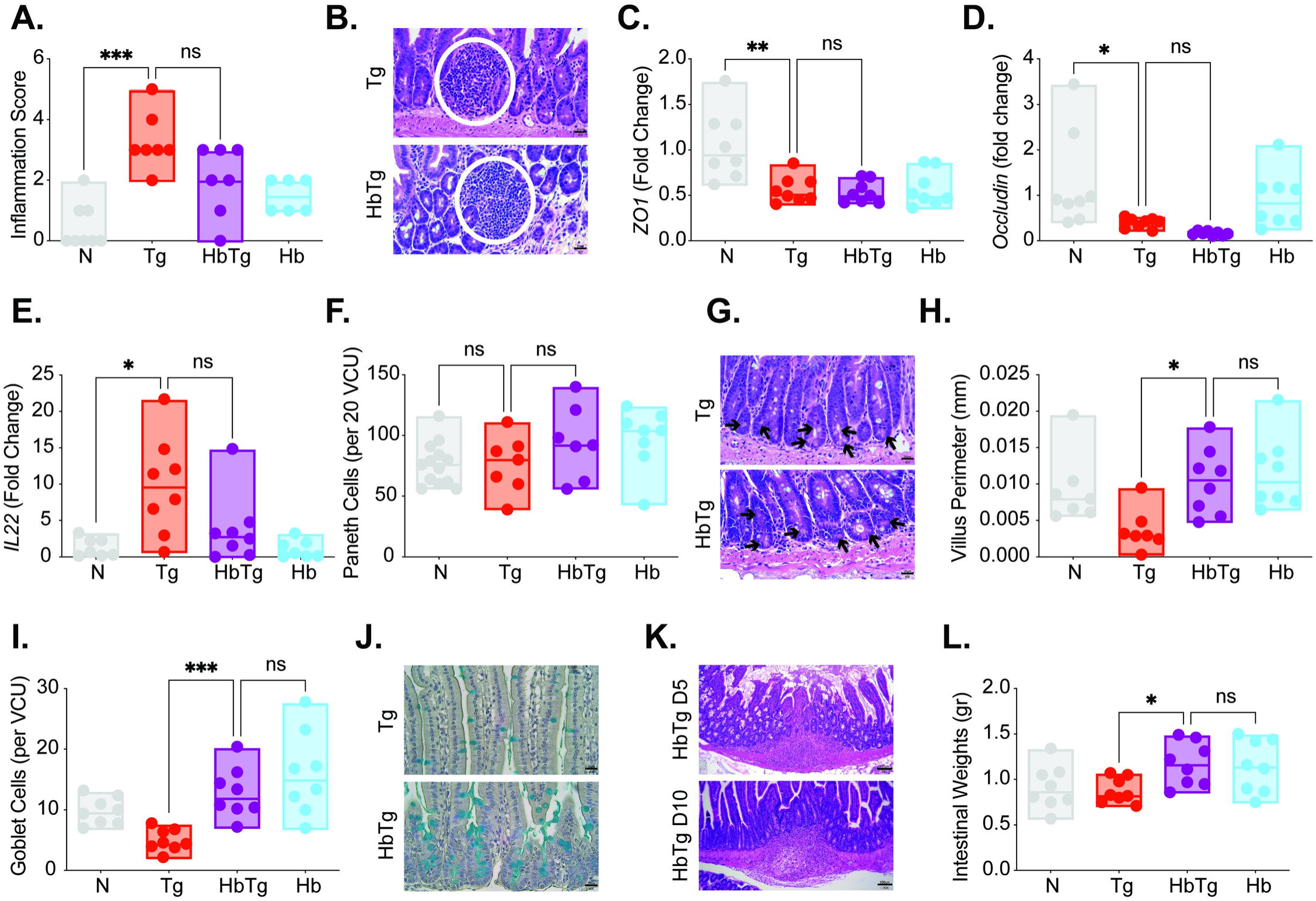
Pathology in the small intestine differs between *T. gondii* and co-infected animals. 200 Hb larvae were given orally to mice 7 days prior to infection with 20 Tg tissue cysts. Small intestines were harvested from mice 5 days post Tg infection, and formalin fixed. 6 μM slides were cut from the paraffin embedded swiss rolls, and stained with hematoxylin and eosin. (A) Average small intestine inflammation pathology score for each mouse, euthanized at 5 post Tg infection. (B) Representative images from Tg and HbTg animals 5 post Tg infection. Scale: 20 μm. White circles depict leukocyte infiltrates (Tg pathology). Gene expression was measured by quantitative RT-PCR at 5 days post-Tg-infection for *Zo1* (C), *Occludin* (D) and *Il-22* (E) in SI2 of the SI. (F) Average number of paneth cells identified on H&E stained sections in 20 villi/crypt units of each experimental group on day 5 post Tg infection. (G) Representative images from Tg and HbTg animals 5 days post Tg infection. Scale: 20 μm. Black arrows depict paneth cells. (H) Average villi perimeter in the proximal small intestine of each experimental group on day 5 post Tg infection. The perimeter of 5 intact continuous villi was measured and averaged. (I) Paraffin embedded swiss rolls were stained with Alcian blue to identify goblet cells. Average number of goblet cells in 5 intact and continuous villi within the proximal small intestine of each experimental group on day 5 post Tg infection. (J) Representative images from Tg and HbTg animals 5 days post Tg infection. Scale: 20 μm. Blue cells depict goblet cells. (K) Representative images of a granuloma in HbTg mice at days 5 and 10 post Tg infection. Scale: 100μm. (L) Small intestines were dissected and weighed. N= 2-5 mice per group per experiment, 2 independent experiments. Normality test followed by ANOVA/Kriskal-Wallis with post-tests were performed for pooled data, n.s. = non significant, * = p<0.05, ** = p<0.01, *** = p<0.001.

After having replicated in the intestinal tissue, Tg rapidly disseminates to other organs, including the spleen and liver. Analysis of spleen and liver sections indicate that pathology of both organs increased over time but that Tg and HbTg infected animals had similar severe pathologies, while Hb infected animals were far less affected (S6 Fig).

## Discussion

Our data confirm the recently published work demonstrating increased mortality in animals coinfected with Tg and Hb compared to Tg alone (28). At first, this may appear counterintuitive. The immunoregulatory effects of helminths have been intensely studied (83–87) and HbTg co-infection was previously shown to improve survival in association with a decrease in serum IFNγ levels (26). Early IFNγ production plays a key role in controlling *T. gondii* replication (88) but IL-10 is also necessary to limit pathology stimulated by the strong inflammatory response (61). The balance of these two cytokines is key to infection outcome (89). Surprisingly, we found very few differences in *Ifnγ* and *Il-10* gene expression between HbTg and Tg animals, despite their survival differences. Tg parasite burden was linked to *Ifnγ* levels: in tissues with high parasite numbers, we recorded high levels of *Ifnγ* (Fig 2). However, compared to Tg infected animals, HpTg infected animals only had decreased levels of MLN *Ifnγ* and SI2 *Il-10* at day 5 post Tg infection, despite increased *Il-4* and *Il-13* in the SI, PP, MLN and SPL. The local decrease in MLN *Ifnγ* was associated with increased MLN Tg burden 5 days later, highlighting the different responses in different tissues.

IFNγ production by CD4^+^ and CD8^+^ T cells is crucial for protective immunity and overall survival of Tg infected animals (90–92). We (Fig 3 and 5A-C) and others (26,28,32) have shown that HbTg co-infection results in decreased IFN γ−producing CD4^+^ and CD8^+^ T cells. NK cells are also critical for driving an effective innate response (49,64,93), including the development of local IL-12 producing DCs and macrophages, that drive effector responses (32,65,94) and both NKT (95) and γδ T cells (68) have been implicated in early response to Tg-induced intestinal pathology. A recent study using a novel IFN γ reporter mouse, demonstrated that CD4 ^+^ T, CD8 ^+^ T, γδ T and NK cells all express IFN γ during Tg infection (96). They found that although IFN γ-producing CD4 ^+^ T and CD8 ^+^ T cells were predominantly responsible for combating acute protection, IFNγ-producing NK and γδ T cell also likely played a role. Our results demonstrate that HpTg co-infection leads to a decrease in all IFN γ−producing lymphocytes (CD8^+^ and CD4^+^ T as well as NK, NKT, γδ T cells) in the MLN as a proportion (Fig 3) and in absolute numbers (S5 F-J Fig), which helps explain the subsequent increase in Tg burden in this organ.

Despite the ablation of the IFN γ responses in all lymphocytes in the MLN of HbTg infected animals 5 days post infection, GZMB production did not follow the same pattern (Fig 4, 5D-I and S5 K-O Fig). GZMB is a molecule involved in CD8 ^+^ T cell and NK cell cytotoxicity (97,98) and we were surprised it was not decreased, like IFN γ, in HbTg animals or associated with better control of Tg parasite loads. However, this may be because Tg itself can inhibit lymphocyte degranulation (99) and GZMB-mediated apoptosis of infected cells (70,100), or the fact that the impact of worm infection on GZMB is unclear (101). Studies have reported that IL-4 production results in increased cytotoxicity in splenic NK cells in the context of *Nippostrongylus brasiliensis* (parasitic nematode) infection (102). Others have found that IL-4 suppresses NK cell cytotoxicity *in vitro* (103–105). GZMB also inhibits tissue repair at mucosal surfaces and is being studied as a therapeutic target for wound healing (106,107). Elevated levels in HbTg compared to Tg animals may be contributing to tissue damage.

Beside the lack of local IFN γ producers, the observed survival differences were also linked to differences in intestinal pathology. Hb and Tg share the small intestinal niche. Tg is known for causing ileitis (inflammation of the ileum, (108). However, pathology in the duodenum and jejenum, both niches for Hb, have also been described (80,109). When mice were co-infected with Tg, Hb was transitioning from a tissue dwelling to a lumen dwelling phase, resulting in pathology in the proximal small intestine. The pathology stemmed from immune effects on the intestinal epithelium (change in secretory cell populations like goblet cells), the granulomas created from an influx of immune cells to the site of infection as well as the damage caused by the worms exiting the intestinal tissue itself. Unsurprisingly we observed that HbTg infected animals had transient increases in goblet cells, *Il-4* and *Il-13* gene expression and a sustained presence of Th2 granulomas compared to Tg infected animals, all associated with Hb infection. We also observed a transient decrease in intestinal permeability and late increases in *Ifnγ* and *Il-10* expression compared to naïve animals (Fig 6). Th2 granulomas are composed mainly of M2 macrophages and eosinophils that promote fibrotic lesions (13). The combination of these granulomas as well as intestinal inflammation stimulated by Tg likely led to intestinal dysfunction and contributed to the increased mortality of the HbTg infected animals. In support of this, we found no pathological differences between the HpTg and Tg groups within the spleen or liver (S6 Fig) or the Tg burdens in these 3 organs (Fig 1). In both Tg and HbTg infected mice, we observed severe pathology by 10 days post Tg infection (S6 Fig).

Previous studies have observed that animals coinfected with Hb and an inflammatory intestinal pathogen have increased mortality associated with significant weight loss approximately 10 days post-infection onwards. These include animals co-infected with Hb and Tg (28), Hb and *Citrobacter rodentium* (110) and Hb and West Nile Virus (WNV) (111). The weight loss has been directly linked to IL-4 mediated events since treating animals with IL-4c had the same effect as an Hb infection (111) and the weight loss, and increased mortality, was reversed in STAT6 knock out animals (110,111). In the case of HbWNV co-infected animals, activation of a succinate-tuft cell-IL-25-IL-4R-intestinal circuit was found to be responsible for the increased mortality and weight loss (111). However, it was linked to intestinal pathology (reduced villus height and bacterial translocation, a read out for intestinal permeability) which we did not observe in our HbTg infected animals.

Others have found that the mechanisms underlying the weight loss observed in HpTg infected animals is unclear, and hypothesised that the increased mortality may be due to tissue stress and proinflammatory responses in certain organs (28). The authors concluded that uncontrolled parasite replication and/or impaired T cell responses were not responsible, but that decreased food intake, poor nutrient absorption, or increased energy expenditure could be responsible for increased mortality in coinfected mice. We agree with this hypothesis. We found few differences in parasite replication and/or impaired responses despite studying two infection time points and 5 different organs (Fig 1-5). We did not measure weight loss past 10 days post infection due to the high mortality in the HbTg infected group. However, infection with Hb and *N. brasiliensis* reduces glucose absorption by the small intestine in a STAT6 dependent manner (66, 67). And most recently, excretory/secretory products from the helminth *Schistosoma mansoni* were shown to inhibit food intake and decrease body fat mass in a STAT6-independent mechanism in animals fed a high fat diet (112).

Our results provide insights into how co-infection with parasites stimulating different arms of the immune system can lead to drastic changes in infection dynamics. Understanding the intricacies of immune responses during co-infections is key to developing models that will provide better translatability to real world situations, where invariably humans, livestock, companion animals and wildlife will be harbouring more than one type of infection.

## Figure legends

**S1 Fig. The spleen pathology scale.** (A). Follicle damage: All follicles are distinct and defined in shape (score of 1, top left). A majority of follicles are distinct and defined in shape (score of 2, top middle). Approximately equal numbers of intact and damaged follicles. (score of 3, top right). The majority of follicles are damaged, with some defined follicles observed (score of 4, bottom left). All follicles are damaged (score of 5, bottom right). Scale 10um. (B) Marginal zone thickness: More than 50% of the follicles have a thick marginal zone (score of 1, left). More than 50% of the follicles have a thin marginal zone (score of 2, middle) More than 50% of the follicles do not have an observable marginal zone (score of 3, right). (C) Size of the light zone of the germinal centre: More than 50% of the follicles have a small germinal center (score of 1, left). More than 50% of the follicles have a large germinal center (score of 2, middle). More than 50% of the follicles have no observable germinal center (score of 3, right).

**S2 Fig. The liver pathology scale.** (A). Necrosis: absence (score of 1, left) and presence (score of 2, right) of necrosis. Scale 10um. (B) Infiltrate Size: All infiltrates observed are organized into small groups (score of 1, left), 75% of infiltrates observed are organized into small groups and 25% into large groups (score of 2, middle left), 50/50 split between small and large infiltrate groups (score of 3, middle right), 100% of infiltrates are organized into large groups (score of 4, right). (C). Proportion of infiltrates: very few infiltrate groups are observed (score of 1, top left), some groups of infiltrates are observed (score of 2, top middle), a medium number of infiltrates is observed (score of 3, top right), many infiltrate groups are observed and occupy the majority of the tissue (score of 4, bottom left), infiltrate groups are distinguishable, and occupy most of the tissue (score of 5, bottom middle), and Infiltrate groups are indistinguishable and occupy the entire tissue (score of 6, bottom right). (D) Perivascular infiltrates: Very few infiltrating leukocytes can be observed in the vessels (score of 1, left). Infiltrating leukocytes are observed in small numbers leaving most of the vessels (score of 2, left middle). Most of the vessels are surrounded by infiltrating leukocytes (score of 3, right middle). All of the vessels are filled with infiltrating leukocytes (score of 4, right).

S3 Fig. **Flow cytometry gating strategy**. Doublets were removed using the FSC-A and FSC-H parameters. Live cells were selected, followed by CD45^+^ cells. The CD3-subset was used to identify NK cells using NK1.1. The CD3^+^ subset was divided into γδ^+^ cells, NKT cells (γδ-αβ^+^NK1.1^+^) and the αβ cells were subdivided into CD4^+^ and CD8^+^ cells.

S4 Fig. **IFN**γ **serum protein levels do not differ between Tg and HbTg infected animals**. IFNγ protein levels were measured in the serum by ELISA at 5 and 10 days post Tg infection. N= 2-4 mice per group per experiment, a minimum of 2 independent experiments. Data was tested for normality (Anderson-Darling test). ANOVA or Kruskal-Wallis tests were performed on parametric/non-parametric pooled data, and when significant, Sidak’s/Dunn’s Multiple comparisons were performed on Hb vs. HbTg and Tg vs. HbTg; ****= *p*<0.0001.

S5 Fig. **Co-infected mice have reduced numbers of MLN IFN**γ **and increased GZMB-producing cells.** 200 Hb larvae were given orally to mice 7 days prior to infection with 20 Tgtissue cysts. (A-E) The cell number of (A) NK (IFNγ+NK1.1^+^CD3-), (B) CD8 T (IFNγ+CD3^+^CD8^+^), (C) NKT (IFNγ+NK1.1^+^CD3^+^), (D) γδ T (IFNγ+ CD3^+^γδ+) and (E) CD4 T cells (IFNγ+CD3^+^CD4^+^) in Tg and HbTg animals. (F-J) The cell number of IFNγ-producing (F) NK (IFNγ+NK1.1^+^CD3-), (G) CD8 T (IFNγ+CD3^+^CD8^+^), (H) NKT (IFNγ+NK1.1^+^CD3^+^), (I) γδ T (IFNγ+ CD3^+^γδ+) and (J) CD4 T cells (IFNγ+CD3^+^CD4^+^) in Tg and HbTg animals. (K-O) The cell number of GZMB-producing (K) NK (GZMB^+^NK1.1^+^CD3-), (L) CD8 T (GZMB^+^CD3^+^CD8^+^), (M) NKT (GZMB+NK1.1^+^CD3^+^), (N) γδ T (GZMB+ CD3^+^γδ+) and (O) CD4 T cells (GZMB+CD3^+^CD4^+^) in Tg and HbTg animals. N= 2-4 mice per group per experiment, 2 independent experiments. Data was tested for normality (Anderson-Darling test). ANOVA or Kruskal-Wallis tests were performed on parametric/non-parametric pooled data including N/Tg/HbTg/Hb groups, and when significant, Sidak’s/Dunn’s Multiple comparisons were performed on Hb vs. HbTg and Tg vs. HbTg; n.s. = non significant, *= *p*<0.05 and ** = *p*<0.01.

S6 Fig. **Pathology in the spleen and liver are similar between *T. gondii* and co-infected animals.** 200 Hb larvae were given orally to mice 7 days prior to infection with 20 Tgtissue cysts. Spleens and livers were harvested from mice 10 days post Tg infection, and formalin fixed. 6 μM slides were cut from the paraffin embedded swiss rolls, and stained with hematoxylin and eosin. Average spleen (left) and liver (right) pathology score for each mouse, euthanized at 10 days post Tg infection. N>3 mice per group per experiment, 2 independent experiments. Data was tested for normality (Anderson-Darling test). ANOVA or Kruskal-Wallis tests were performed on parametric/non-parametric pooled data including N/Tg/HbTg/Hb groups, and when significant, Sidak’s/Dunn’s Multiple comparisons were performed on N vs Tg and Tg vs. HbTg; n.s. = non significant, * = *p*<0.05 and ** = *p*<0.01.

S1 Table 1. **Small intestine pathology scales.** The total score for each animal was calculated by adding the score for the pathology associated with epithelial integrity and inflammatory cell infiltrate.

S2 Table 2. **Spleen and Liver Pathology Scales**. For the spleen, the total score for each animal was calculated by adding the score for the pathology associated with the follicle shape, the marginal zone and the germinal centre. For the liver, the total score for each animal was calculated by adding the score for the pathology associated with necrosis, the infiltrate size, the proportion of infiltrates and the nature of the perivascular infiltrates.

## ACKNOWLEDGEMENTS

We would like to thank the University of Calgary LESARC Facility technicians, especially Dawn Martin for their continued support as well as the University of Calgary’s Diagnostic Services Unit, especially Sue Calder-Lodge and JJ Larios for their patience and expertise.

This works was funded through Dr. Finney’s grants from the Canadian Foundation for Innovation and the Natural Sciences and Engineering Research Council of Canada (NSERC), as well as scholarships for Dr Anupama Ariyaratne (NSERC Create in Host Parasite Interactions), Namratha Badawadagi (University of Calgary Markin scholarship), Kayla Bailey and Emma Forrester (University of Calgary PURE Scholarship), Dr Joel Bowron and Beverly Dong (NSERC), Camila Gaio and Manfred Ritz (Mitacs Globalinks Scholarships), Shashini Perera (Alberta Graduate Excellence Scholarship) and Dr Edina K Szabo (UCalgary Eyes High Postdoctoral Scholarship).

## DATA AVAILABILITY

All relevant data are within the manuscript and its Supporting Information files

## Notes

### Competing Interest Statement

The authors have declared no competing interest.

### Summary of Updates

All figures were revised for clarity and additional data was added for completeness.

## References

1. Hoarau AOG, Mavingui P, Lebarbenchon C. Coinfections in wildlife: Focus on a neglected aspect of infectious disease epidemiology. PLoS Pathog. 2020 Sep;16(9):e1008790-.

2. Shen SS, Qu XY, Zhang WZ, Li J, Lv ZY. Infection against infection: parasite antagonism against parasites, viruses and bacteria. Infect Dis Poverty. 2019;8(1):49.

3. Graham AL, Cattadori IM, Lloyd-Smith JO, Ferrari MJ, Bjørnstad ON. Transmission consequences of coinfection: cytokines writ large? Vol. 23, Trends in Parasitology. 2007. p. 284–91.

4. Brunton L, Hilal-Dandan R, Knollmann B. Goodman and Gilman’s The Pharmacological Basis of Therapeutics, 13th Edition. 13th ed. USA: McGraw-Hill Education; 2018.

5. Avramenko RW, Bras A, Redman EM, Woodbury MR, Wagner B, Shury T, et al. High species diversity of trichostrongyle parasite communities within and between Western Canadian commercial and conservation bison herds revealed by nemabiome metabarcoding. Parasit Vectors. 2018;11(1):299.

6. Clerc M, Devevey G, Fenton A, Pedersen AB. Antibodies and coinfection drive variation in nematode burdens in wild mice. Int J Parasitol. 2018;48(9–10):785–92.

7. Cooper PJ, Chico M, Sandoval C, Espinel I, Guevara A, Levine MM, et al. Human infection with Ascaris lumbricoides is associated with suppression of the interleukin-2 response to recombinant cholera toxin B subunit following vaccination with the live oral cholera vaccine CVD 103-HgR. Infect Immun. 2001;69(3):1574–80.

8. Graham AL, Cattadori IM, Lloyd-Smith JO, Ferrari MJ, Bjørnstad ON. Transmission consequences of coinfection: cytokines writ large? Vol. 23, Trends in Parasitology. 2007. p. 284–91.

9. Su Z, Segura M, Stevenson MM. Reduced protective efficacy of a blood-stage malaria vaccine by concurrent nematode infection. Infect Immun. 2006 Apr;74(4):2138–44.

10. Koboziev I, Karlsson F, Grisham MB. Gut-associated lymphoid tissue, T cell trafficking, and chronic intestinal inflammation. Vol. 1207, Annals of the New York Academy of Sciences. Blackwell Publishing Inc.; 2010.

11. Stevens L, Martinez-Ugalde I, King E, Wagah M, Absolon D, Bancroft R, et al. Ancient diversity in host-parasite interaction genes in a model parasitic nematode. bioRxiv [Internet]. 2023 Apr 17 [cited 2023 Sep 18];2023.04.17.535870. Available from: https://www.biorxiv.org/content/10.1101/2023.04.17.535870v1

12. Reynolds L a., Filbey KJ, Maizels RM. Immunity to the model intestinal helminth parasite Heligmosomoides polygyrus. Semin Immunopathol. 2012 Nov;34(6):829–46.

13. Ariyaratne A, Finney CAM. Eosinophils and macrophages within the Th2-induced granuloma: Balancing killing and healing in a tight space. Infect Immun. 2019 Jul;87(10):e00127–19.

14. Smith KA, Filbey KJ, Reynolds LA, Hewitson JP, Harcus Y, Boon L, et al. Low-level regulatory T-cell activity is essential for functional type-2 effector immunity to expel gastrointestinal helminths. Mucosal Immunol. 2016;9(2):428–43.

15. Filbey KJ, Grainger JR, Smith KA, Boon L, van Rooijen N, Harcus Y, et al. Innate and adaptive type 2 immune cell responses in genetically controlled resistance to intestinal helminth infection. Immunol Cell Biol. 2014;92(5):436–48.

16. Gregg B, Taylor BC, John B, Tait-Wojno ED, Girgis NM, Miller N, et al. Replication and distribution of toxoplasma gondii in the small intestine after oral infection with tissue cysts. Infect Immun. 2013 May;81(5):1635–43.

17. Denkers EY, Icardo R, Gazzinelli T, Martin D, Sher A. Emergence of NKI.1 + Cells as Effectors of IFN-3’ Dependent Immunity to Toxoplasma gondii in MHC Class I-deficient Mice. Journal of Experimental Medicine. 1993;178(5):1465–72.

18. Sher A, Oswald IP, Hieny S, Gazzinelli RT. Toxoplasma gondii Induces a T-Independent IFN-)/ Response in Natural Killer Cells That Requires Both Adherent Accessory Cells and Tumor Necrosis Factor-a. The journal of Immunology. 1993;150(9):3982–9.

19. Goldszmid RS, Caspar P, Rivollier A, White S, Dzutsev A, Hieny S, et al. NK Cell-Derived Interferon-γ Orchestrates Cellular Dynamics and the Differentiation of Monocytes into Dendritic Cells at the Site of Infection. Immunity. 2012 Jun;36(6):1047– 59.

20. Combe CL, Curiel TJ, Moretto MM, Khan IA. NK Cells Help To Induce CD8 LJ -T-Cell Immunity against Toxoplasma gondii in the Absence of CD4 LJ T Cells. Society. 2005 Aug;73(8):4913–21.

21. Nakano Y, Hisaeda H, Sakai T, Zhang M, Maekawa Y, Zhang T, et al. Granule-dependent killing of Toxoplasma gondii by CD8 + T cells. Immunology. 2001;104(3):289–98.

22. Kasper LH, Matsuura T, Fonseka S, Arruda J, Channon JY, Khan IA. Induction of gammadelta T cells during acute murine infection with Toxoplasma gondii. The Journal of Immunology. 1996 Dec;157(12):5521.

23. Ronet C, Darche S, de Moraes ML, Miyake S, Yamamura T, Louis JA, et al. NKT Cells Are Critical for the Initiation of an Inflammatory Bowel Response against Toxoplasma gondii. The Journal of Immunology. 2005 Jul;175(2):899–908.

24. Nakano Y, Hisaeda H, Sakai T, Ishikawa H, Zhang M, Maekawa Y, et al. Roles of NKT cells in resistance against infection with Toxoplasma gondii and in expression of heat shock protein 65 in the host macrophages. Microbes Infect. 2002;4(1):1–11.

25. Ahmed N, French T, Rausch S, Kühl A, Hemminger K, Dunay IR, et al. Toxoplasma Co-infection Prevents Th2 Differentiation and Leads to a Helminth-Specific Th1 Response. Front Cell Infect Microbiol. 2017;7(July):1–12.

26. Khan IA, Hakak R, Eberle K, Sayles P, Weiss LM, Urban JF. Coinfection with Heligmosomoides polygyrus fails to establish CD8 + T-cell immunity against Toxoplasma gondii. Infect Immun. 2008;76(3):1305–13.

27. Marple A, Wu W, Shah S, Zhao Y, Du P, Gause WC, et al. Cutting Edge: Helminth Coinfection Blocks Effector Differentiation of CD8 T Cells through Alternate Host Th2- and IL-10–Mediated Responses. The Journal of Immunology. 2017 Jan;198(2):634–9.

28. Rovira-Diaz E, El-Naccache DW, Reyes J, Zhao Y, Nasuhidehnavi A, Chen F, et al. The Impact of Helminth Coinfection on Innate and Adaptive Immune Resistance and Disease Tolerance during Toxoplasmosis. The Journal of Immunology [Internet]. 2022 Dec 1 [cited 2023 Jun 8];209(11):2160–71. Available from: 10.4049/jimmunol.2200504

29. Perona-Wright G, Mohrs K, Szaba FM, Kummer LW, Madan R, Karp CL, et al. Systemic but Not Local Infections Elicit Immunosuppressive IL-10 Production by Natural Killer Cells. Cell Host Microbe. 2009 Dec;6(6):503–12.

30. Liesenfeld O, Dunay IR, Erb KJ. Infection with Toxoplasma gondii reduces established and developing Th2 responses induced by Nippostrongylus brasiliensis infection. Infect Immun. 2004 Jul;72(7):3812–22.

31. Coomes SM, Pelly VS, Kannan Y, Okoye IS, Czieso S, Entwistle LJ, et al. IFNγ and IL-12 Restrict Th2 Responses during Helminth/Plasmodium Co-Infection and Promote IFNγ from Th2 Cells. PLoS Pathog. 2015 Jul;11(7).

32. Marple A, Wu W, Shah S, Zhao Y, Du P, Gause WC, et al. Cutting Edge: Helminth Coinfection Blocks Effector Differentiation of CD8 T Cells through Alternate Host Th2- and IL-10–Mediated Responses. The Journal of Immunology. 2017 Jan;198(2):634–9.

33. Shi Y, Zhang P, Wang G, Liu X, Sun X, Zhang X, et al. Description of organ-specific phenotype, and functional characteristics of tissue resident lymphocytes from liver transplantation donor and research on immune tolerance mechanism of liver. Oncotarget. 2018;9(21):15552–65.

34. Moreau MR, Wijetunge DSS, Bailey ML, Gongati SR, Goodfield LL, Hewage EMKK, et al. Growth in Egg Yolk Enhances Salmonella Enteritidis Colonization and Virulence in a Mouse Model of Human Colitis. PLoS One [Internet]. 2016 Mar 1 [cited 2023 Jun 13];11(3):e0150258. Available from: https://journals.plos.org/plosone/article?id=10.1371/journal.pone.0150258

35. Moreau MR, Wijetunge DSS, Bailey ML, Gongati SR, Goodfield LL, Hewage EMKK, et al. Growth in Egg Yolk Enhances Salmonella Enteritidis Colonization and Virulence in a Mouse Model of Human Colitis. Chang YF, editor. PLoS One. 2016 Mar;11(3):e0150258.

36. Pfaffl MW. Relative quantification. 2004;63–82.

37. Costa JM, Bretagne S. Variation of B1 gene and AF146527 repeat element copy numbers according to Toxoplasma gondii strains assessed using real-time quantitative PCR. J Clin Microbiol. 2012 Apr;50(4):1452–4.

38. Cossarizza A, Chang HD, Radbruch A, Acs A, Adam D, Adam-Klages S, et al. Guidelines for the use of flow cytometry and cell sorting in immunological studies (second edition). Eur J Immunol. 2019;49(10):1457–973.

39. Bryant V. The Life Cycle of *Nematospiroides dubius*, Baylis, 1926 (Nematoda: Heligmosomidae). J Helminthol. 1973 Sep;47(3).

40. Dubey JP, Dubey’ JP. Bradyzoite-induced murine toxoplasmosis: stage conversion, pathogenesis, and tissue cyst formation in mice fed bradyzoites of different strains of Toxoplasma gondii. J Eukaryot Microbiol. 1997 Jan;44(6):592–602.

41. Zenner L, Darcy F, Capron A, Cesbron-Delauw MF. Toxoplasma gondii: Kinetics of the Dissemination in the Host Tissues during the Acute Phase of Infection of Mice and Rats. Exp Parasitol. 1998;90:86–94.

42. Gregg B, Taylor BC, John B, Tait-Wojno ED, Girgis NM, Miller N, et al. Replication and distribution of toxoplasma gondii in the small intestine after oral infection with tissue cysts. Infect Immun. 2013 May;81(5):1635–43.

43. Hunter CA, Subauste CS, Van Cleave VH, Remington JS. Production of gamma interferon by natural killer cells from Toxoplasma gondii-infected SCID mice: regulation by interleukin-10, interleukin-12, and tumor necrosis factor alpha. Infect Immun. 1994 Jul;62(7):2818–24.

44. Khan IA, Thomas SY, Moretto MM, Lee FS, Islam SA, Combe C, et al. CCR5 is essential for NK cell trafficking and host survival following Toxoplasma gondii infection. PLoS Pathog. 2006;2(6):0484–500.

45. Suzuki Y, Orellana M, Schreiber R, Remington J. Interferon-gamma: the major mediator of resistance against Toxoplasma gondii. Science (1979). 1988 Apr;240(4851):516–8.

46. Suzuki Y, Sa Q, Gehman M, Ochiai E. Interferon-gamma- and perforin-mediated immune responses for resistance against Toxoplasma gondii in the brain. Vol. 13, Expert reviews in molecular medicine. 2011.

47. Scharton-Kersten TM, Wynn TA, Denkers EY, Bala S, Grunvald E, Hieny S, et al. In the absence of endogenous IFN-gamma, mice develop unimpaired IL-12 responses to Toxoplasma gondii while failing to control acute infection. The Journal of Immunology. 1996 Nov;157(9):4045.

48. Suzuki Y, Remington JS. against toxoplasmosis in mice. protective effect of Lyt-2+ immune T cells The effect of anti-IFN-gamma antibody on the. The Journal of Immunology. 1990;144:1954–6.

49. Denkers EY, Icardo R, Gazzinelli T, Martin D, Sher A. Emergence of NKI.1 + Cells as Effectors of IFN-3’ Dependent Immunity to Toxoplasma gondii in MHC Class I-deficient Mice. Journal of Experimental Medicine. 1993;178(5):1465–72.

50. Norose K, Mun HS, Aosai F, Chen M, Piao LX, Kobayashi M, et al. IFN-γ-regulated Toxoplasma gondii distribution and load in the murine eye. Invest Ophthalmol Vis Sci. 2003 Oct;44(10):4375–81.

51. Dimier IH. Interferon-y-activated primary enterocytes inhibit Toxoplasma gondii replication: a role for intracellular iron. Immunology. 1998;94:488–95.

52. Speer CA, Dubey JP. Ultrastructure of early stages of infections in mice fed Toxoplasma gondii oocysts. Parasitology. 1998/01/01. 1998;116(1):35–42.

53. Dubey JP, Ferreira LR, Martins J, Mcleod R. Oral oocyst-induced mouse model of toxoplasmosis: Effect of infection with Toxoplasma gondii strains of different genotypes, dose, and mouse strains (transgenic, out-bred, in-bred) on pathogenesis and mortality. Parasitology. 2012 Jan;139(1):1–13.

54. Lewis JW, Bryant V. The distribution of nematospiroides dubiiis within the small intestine of laboratory mice. J Helminthol. 1976 Sep 5;50(3):163–71.

55. Paludan SR, Lovmand J, Ellermann-Eriksen S, Mogensen SC. Effect of IL-4 and IL-13 on IFN-γ -induced production of nitric oxide in mouse macrophages infected with herpes simplex virus type 2. FEBS Lett. 1997 Sep;414(1).

56. Klein SA, Dobmeyer JM, Dobmeyer TS, Pape M, Ottmann OG, Helm EB, et al. Demonstration of the Th1 to Th2 cytokine shift during the course of HIV-1 infection using cytoplasmic cytokine detection on single cell level by flow cytometry. AIDS. 1997;11(9).

57. Wurtz O, Bajénoff M, Guerder S. ILLJ4LJmediated inhibition of IFNLJγ production by CD4+ T cells proceeds by several developmentally regulated mechanisms. Int Immunol. 2004 Mar;16(3):501–8.

58. Mckenzie GJ, Fallon PG, Emson CL, Grencis RK, Mckenzie ANJ. Simultaneous Disruption of Interleukin (IL)-4 and IL-13 Defines Individual Roles in T Helper Cell Type 2-mediated Responses. J Exp Med. 1999;189(10):1565–72.

59. Urban JF, Noben-Trauth N, Donaldson DD, Madden KB, Morris SC, Collins M, et al. IL-13, IL-4R, and Stat6 Are Required for the Expulsion of the Gastrointestinal Nematode Parasite Nippostrongylus brasiliensis either parasite, disruption of the IL-4 gene or inhibition of IL-4 with an antibody that blocks IL-4 receptor chain (IL-4R) funct. Immunity. 1998;8:255–64.

60. Oeser K, Schwartz C, Voehringer D. Conditional IL-4/IL-13-deficient mice reveal a critical role of innate immune cells for protective immunity against gastrointestinal helminths. Mucosal Immunol. 2015 May 8;8(3):672–82.

61. Suzuki Y, Sher A, Yap G, Park D, Neyer LE, Liesenfeld O, et al. IL-10 Is Required for Prevention of Necrosis in the Small Intestine and Mortality in Both Genetically Resistant BALB/c and Susceptible C57BL/6 Mice Following Peroral Infection with Toxoplasma gondii. The Journal of Immunology [Internet]. 2000 May 15 [cited 2020 Jul 15];164(10):5375–82. Available from: http://www.jimmunol.org/content/164/10/5375http://www.jimmunol.org/content/164/10/5375.full#ref-list-1

62. Denkers EY, Sher A, Gazzinelli RT. CD8+ T-cell interactions with Toxoplasma gondii: implications for processing of antigen for class-I-restricted recognition. Res Immunol. 1993;144(1):51–7.

63. Khan IA, Green WR, Kasper LH, Green KA, Schwartzman JD. Immune CD8+ T Cells Prevent Reactivation of Toxoplasma gondii Infection in the Immunocompromised Host. Infect Immun. 1999 Nov;67(11).

64. Sher A, Oswald IP, Hieny S, Gazzinelli RT. Toxoplasma gondii Induces a T-Independent IFN-)/ Response in Natural Killer Cells That Requires Both Adherent Accessory Cells and Tumor Necrosis Factor-a. The journal of Immunology. 1993;150(9):3982–9.

65. Goldszmid RS, Caspar P, Rivollier A, White S, Dzutsev A, Hieny S, et al. NK Cell-Derived Interferon-γ Orchestrates Cellular Dynamics and the Differentiation of Monocytes into Dendritic Cells at the Site of Infection. Immunity. 2012 Jun;36(6):1047– 59.

66. Edelblum KL, Singh G, Odenwald MA, Lingaraju A, El Bissati K, McLeod R, et al. γδ intraepithelial lymphocyte migration limits transepithelial pathogen invasion and systemic disease in mice. Gastroenterology. 2015 Jun;148(7):1417–26.

67. Khan IA, Hwang S, Moretto M. Toxoplasma gondii: CD8 T cells cry for CD4 help. Front Cell Infect Microbiol. 2019;9(MAY).

68. Nakano Y, Hisaeda H, Sakai T, Ishikawa H, Zhang M, Maekawa Y, et al. Roles of NKT cells in resistance against infection with Toxoplasma gondii and in expression of heat shock protein 65 in the host macrophages. Microbes Infect. 2002;4(1):1–11.

69. Bhadra R, Gigley JP, Khan IA. The CD8 T-cell road to immunotherapy of toxoplasmosis. Vol. 3, Immunotherapy. 2011. p. 789–801.

70. Yamada T, Tomita T, Weiss LM, Orlofsky A. Toxoplasma gondii inhibits granzyme B-mediated apoptosis by the inhibition of granzyme B function in host cells. Int J Parasitol. 2011 May;41(6):595–607.

71. Denkers EY, Yap G, Scharton-Kersten T, Charest H, Butcher BA, Caspar P, et al. Perforin-Mediated Cytolysis Plays a Limited Role in Host Resistance to Toxoplasma gondii. Journal of Immunology. 1997;159:1903–4.

72. Fujiwara A, Kawai Y, Sekikawa S, Horii T, Yamada M, Mitsufuji S, et al. Villus epithelial injury induced by infection with the nematode Nippostrongylus brasiliensis is associated with upregution of Granzyme B V. Journal of Parasitology. 2004 Oct;90(5):1019–26.

73. Mosconi I, Dubey LK, Volpe B, Esser-von Bieren J, Zaiss MM, Lebon L, et al. Parasite proximity drives the expansion of regulatory T cells in Peyer’s patches following intestinal helminth infection. Infect Immun. 2015;83(9):3657–65.

74. Mitsunaga T, Norose K, Aosai F, Horie H, Ohnuma N, Yano A. Infection dynamics of Toxoplasma gondii in gut-associated tissues after oral infection: The role of Peyer’s patches. Parasitol Int. 2019;68(1):40–7.

75. Briceño MP, Nascimento LAC, Nogueira NP, Barenco PVC, Ferro EAV, Rezende-Oliveira K, et al. Toxoplasma gondii Infection Promotes Epithelial Barrier Dysfunction of Caco-2 Cells. Journal of Histochemistry and Cytochemistry [Internet]. 2016 Aug 1 [cited 2023 Jun 7];64(8):459. Available from: /pmc/articles/PMC4971781/

76. Ramírez-Flores CJ, Cruz-Mirón R, Lagunas-Cortés N, Mondragón-Castelán M, Mondragon-Gonzalez R, González-Pozos S, et al. Toxoplasma gondii excreted/secreted proteases disrupt intercellular junction proteins in epithelial cell monolayers to facilitate tachyzoites paracellular migration. Cell Microbiol [Internet]. 2021 Mar 1 [cited 2023 Jun 7];23(3):e13283. Available from: https://onlinelibrary.wiley.com/doi/full/10.1111/cmi.13283

77. Couturier-Maillard A, Froux N, Piotet-Morin J, Michaudel C, Brault L, Le Bérichel J, et al. Interleukin-22-deficiency and microbiota contribute to the exacerbation of Toxoplasma gondii-induced intestinal inflammation article. Mucosal Immunol [Internet]. 2018 Jul 1 [cited 2023 Jun 7];11(4):1181–90. Available from: http://www.mucosalimmunology.org/article/S1933021922004755/fulltext

78. Raetz M, Hwang SH, Wilhelm CL, Kirkland D, Benson A, Sturge CR, et al. Parasite-induced TH1 cells and intestinal dysbiosis cooperate in IFN-γ-dependent elimination of Paneth cells. Nat Immunol [Internet]. 2013 Feb [cited 2023 Jun 7];14(2):136–42. Available from: https://pubmed.ncbi.nlm.nih.gov/23263554/

79. Burger E, Araujo A, López-Yglesias A, Rajala MW, Geng L, Levine B, et al. Loss of Paneth cell autophagy causes acute susceptibility to Toxoplasma gondii-mediated inflammation. Cell Host Microbe [Internet]. 2018 Feb 2 [cited 2023 Jun 7];23(2):177. Available from: /pmc/articles/PMC6179445/

80. Trevizan AR, Vicentino-Vieira SL, da Silva Watanabe P, Góis MB, de Melo G de AN, Garcia JL, et al. Kinetics of acute infection with Toxoplasma gondii and histopathological changes in the duodenum of rats. Exp Parasitol. 2016 Jun 1;165:22–9.

81. Specian RD, Oliver MG. Functional biology of intestinal goblet cells. Vol. 260, American Journal of Physiology - Cell Physiology. American Physiological Society Bethesda, MD; 1991.

82. Ariyaratne A, Yong Kim S, J Pollo SM, Perera S, Liu H, T Nguyen WN, et al. Trickle infection with Heligmosomoides polygyrus results in decreased worm burdens but increased intestinal inflammation and scarring OPEN ACCESS EDITED BY. Front Immunol [Internet]. 2022 [cited 2023 Jun 7];13:1020056. Available from: https://or.ucr.

83. Finney CAM, Taylor MD, Wilson MS, Maizels RM. Expansion and activation of CD4+CD25+regulatory T cells in Heligmosomoides polygyrus infection. Eur J Immunol. 2007;37(7):1874–86.

84. Wilson MS, Taylor MD, Balic A, Finney CAM, Lamb JR, Maizels RM. Suppression of allergic airway inflammation by helminth-induced regulatory T cells. Journal of Experimental Medicine. 2005;202(9).

85. Grainger JR, Smith KA, Hewitson JP, McSorley HJ, Harcus Y, Filbey KJ, et al. Helminth secretions induce de novo T cell Foxp3 expression and regulatory function through the TGF-β pathway. J Exp Med. 2010;207(11):2331–41.

86. Johnston CJC, Smyth DJ, Kodali RB, White MPJ, Harcus Y, Filbey KJ, et al. A structurally distinct TGF-β mimic from an intestinal helminth parasite potently induces regulatory T cells. Nature Communications 2017 8:1 [Internet]. 2017 Nov 23 [cited 2021 Sep 6];8(1):1–13. Available from: https://www.nature.com/articles/s41467-017-01886-6

87. Osbourn M, Soares DC, Vacca F, Cohen ES, Scott IC, Gregory WF, et al. HpARI Protein Secreted by a Helminth Parasite Suppresses Interleukin-33. Immunity. 2017;47(4):739–751.e5.

88. Scharton-Kersten TM, Wynn TA, Denkers EY, Bala S, Grunvald E, Hieny S, et al. In the absence of endogenous IFN-gamma, mice develop unimpaired IL-12 responses to Toxoplasma gondii while failing to control acute infection. The Journal of Immunology. 1996 Nov;157(9):4045.

89. Perona-Wright G, Mohrs K, Szaba FM, Kummer LW, Madan R, Karp CL, et al. Systemic but Not Local Infections Elicit Immunosuppressive IL-10 Production by Natural Killer Cells. Cell Host Microbe. 2009 Dec;6(6):503–12.

90. Dubey JP. Infectivity and Pathogenicity of Toxoplasma gondii Oocysts for Cats. J Parasitol. 1996 Dec;82(6):957–61.

91. Plonquet A, Bassez G, Authier FJ, Dray JM, Farcet JP, Gherardi RK. Toxoplasmic myositis as a presenting manifestation of idiopathic CD4 lymphocytopenia. Muscle Nerve. 2003 Jun;27(6):761–5.

92. Suzuki Y, Orellana M, Schreiber R, Remington J. Interferon-gamma: the major mediator of resistance against Toxoplasma gondii. Science (1979). 1988 Apr;240(4851):516–8.

93. Hunter CA, Subauste CS, Van Cleave VH, Remington JS. Production of gamma interferon by natural killer cells from Toxoplasma gondii-infected SCID mice: regulation by interleukin-10, interleukin-12, and tumor necrosis factor alpha. Infect Immun. 1994 Jul;62(7):2818–24.

94. Wilson DC, Matthews S, Yap GS. IL-12 Signaling Drives CD8 + T Cell IFN-γ Production and Differentiation of KLRG1 + Effector Subpopulations during Toxoplasma gondii Infection. The Journal of Immunology. 2008 May;180(9):5935–45.

95. Ronet C, Darche S, de Moraes ML, Miyake S, Yamamura T, Louis JA, et al. NKT Cells Are Critical for the Initiation of an Inflammatory Bowel Response against Toxoplasma gondii. The Journal of Immunology. 2005 Jul;175(2):899–908.

96. Nishiyama S, Pradipta A, Ma JS, Sasai M, Yamamoto M. T cell-derived interferon-γ is required for host defense to Toxoplasma gondii. Parasitol Int. 2020 Apr 1;75:102049.

97. Dotiwala F, Mulik S, Polidoro RB, Ansara JA, Burleigh BA, Walch M, et al. Killer lymphocytes use granulysin, perforin and granzymes to kill intracellular parasites. Nat Med. 2016 Feb;22(2):210–6.

98. Nakano Y, Hisaeda H, Sakai T, Zhang M, Maekawa Y, Zhang T, et al. Granule-dependent killing of Toxoplasma gondii by CD8 + T cells. Immunology. 2001;104(3):289–98.

99. Bhandage AK, Friedrich LM, Kanatani S, Jakobsson-Björkén S, Escrig-Larena JI, Wagner AK, et al. GABAergic signaling in human and murine NK cells upon challenge with Toxoplasma gondii. J Leukoc Biol [Internet]. 2021 Oct 1 [cited 2023 Sep 15];110(4):617–28. Available from: https://pubmed.ncbi.nlm.nih.gov/34028876/

100. Nash P, Purner M, Leon R, Clarke P, Duke R, Curiel T. Toxoplasma gondii-Infected Cells Are Resistant to Multiple Inducers of Apoptosis. The Journal of Immunology. 1998 Feb;160(4):1824–30.

101. Fujiwara A, Kawai Y, Sekikawa S, Horii T, Yamada M, Mitsufuji S, et al. Villus epithelial injury induced by infection with the nematode Nippostrongylus brasiliensis is associated with upregution of Granzyme B V. Journal of Parasitology. 2004 Oct;90(5):1019–26.

102. Kiniwa T, Enomoto Y, Terazawa N, Omi A, Miyata N, Ishiwata K, et al. NK cells activated by Interleukin-4 in cooperation with Interleukin-15 exhibit distinctive characteristics. Proceedings of the National Academy of Sciences. 2016 Sep;113(36):10139–44.

103. Brady J, Carotta S, Thong RPL, Chan CJ, Hayakawa Y, Smyth MJ, et al. The Interactions of Multiple Cytokines Control NK Cell Maturation. The Journal of Immunology. 2010 Dec;185(11):6679–88.

104. Bratke K, Goettsching H, Kuepper M, Geyer S, Luttmann W, Virchow JC. Interleukin-4 suppresses the cytotoxic potential of in vitro generated, adaptive regulatory CD4+ T cells by down-regulation of granzyme B. Immunology. 2009 Jul;127(3):338–44.

105. Ohayon D, Krishnamurthy D, Brusilovsky M, Waggoner S. IL-4 and IL-13 modulate natural killer cell responses under inflammatory conditions. The Journal of Immunology. 2017 May;198(11):194.

106. Turner CT, Hiroyasu S, Granville DJ. Granzyme B as a therapeutic target for wound healing. Expert Opin Ther Targets [Internet]. 2019 Sep 2 [cited 2023 Sep 15];23(9):745–54. Available from: https://pubmed.ncbi.nlm.nih.gov/31461387/

107. Lim YS, Lee AG, Jiang X, Scott JM, Cofie A, Kumar S, et al. NK cell-derived extracellular granzyme B drives epithelial ulceration during HSV-2 genital infection. Cell Rep [Internet]. 2023 Apr 25 [cited 2023 Sep 18];42(4). Available from: https://pubmed.ncbi.nlm.nih.gov/37071533/

108. Liesenfeld O. Oral infection of C57BL/6 mice with Toxoplasma gondii: a new model of inflammatory bowel disease? J Infect Dis [Internet]. 2002 Feb 15 [cited 2023 Jun 12];185 Suppl 1(SUPPL. 1). Available from: https://pubmed.ncbi.nlm.nih.gov/11865446/

109. Vicentino-Vieira SL, Góis MB, Trevizan AR, de Lima LL, Leatte EP, Nogueira de Melo G de A, et al. Toxoplasma gondii infection causes structural changes in the jejunum of rats infected with different inoculum doses. Life Sci. 2017 Dec 15;191:141–9.

110. Chen CC, Louie S, McCormick B, Walker WA, Shi HN. Concurrent infection with an intestinal helminth parasite impairs host resistance to enteric Citrobacter rodentium and enhances Citrobacter-induced colitis in mice. Infect Immun. 2005 Sep;73(9):5468–81.

111. Desai P, Janova H, White JP, Stappenbeck TS, Thackray LB, Diamond Correspondence MS, et al. Enteric helminth coinfection enhances host susceptibility to neurotropic flaviviruses via a tuft cell-IL-4 receptor signaling axis. Cell [Internet]. 2021 [cited 2023 Jun 7];184. Available from: 10.1016/j.cell.2021.01.051

112. Zande HJP van der, Gonzalez MA, Ruiter K de, Wilbers R, Garcia-Tardón N, Huizen M van, et al. The helminth glycoprotein omega-1 improves metabolic homeostasis in obese mice through type-2 immunity-independent inhibition of food intake. bioRxiv [Internet]. 2020 Jul 3 [cited 2023 Jun 13];2020.07.03.186254. Available from: https://www.biorxiv.org/content/10.1101/2020.07.03.186254v1

