## Supplementary figures and images for "*Heligmosomoides bakeri* and *Toxoplasma gondii* co-infection leads to increased mortality associated with intestinal pathology"

### Supplemental Figure 1

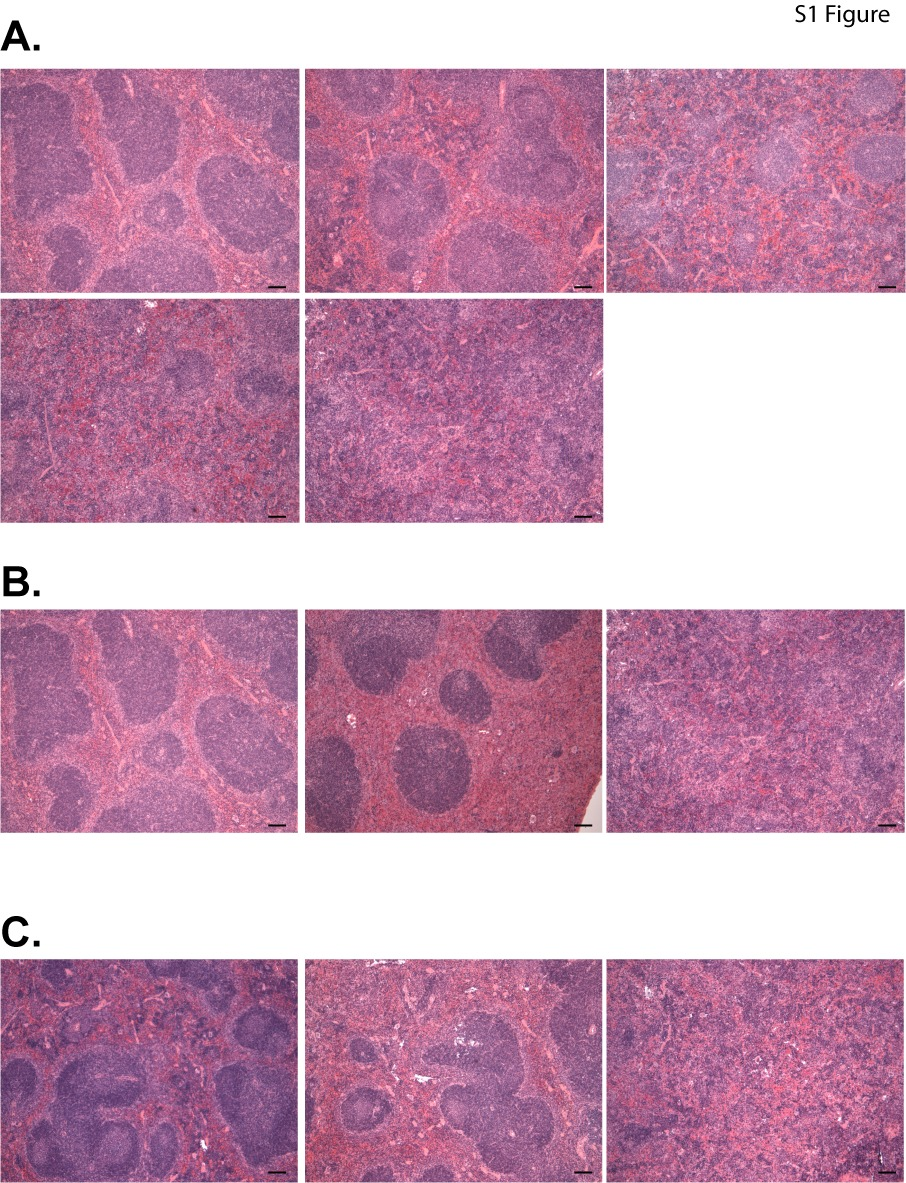

### Supplemental Figure 2

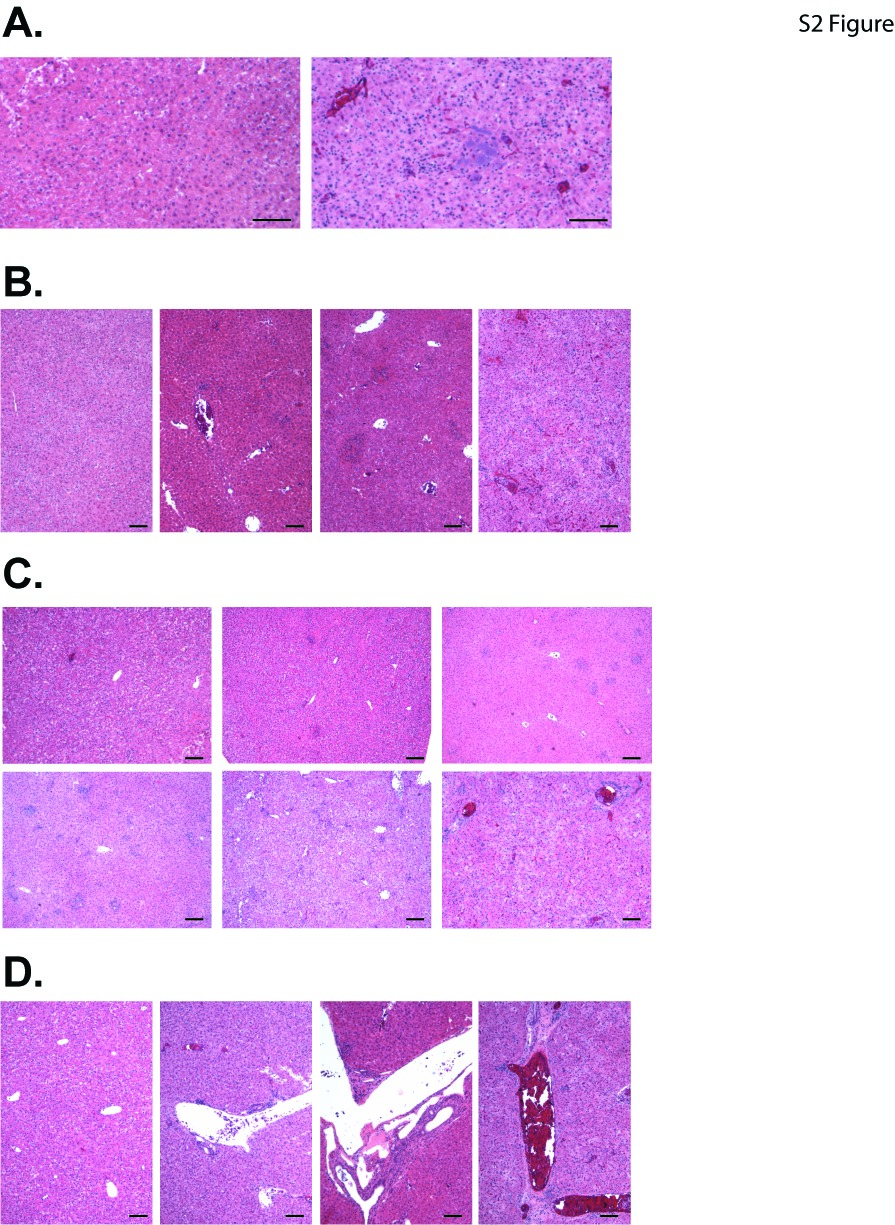

### Supplemental Figure 3

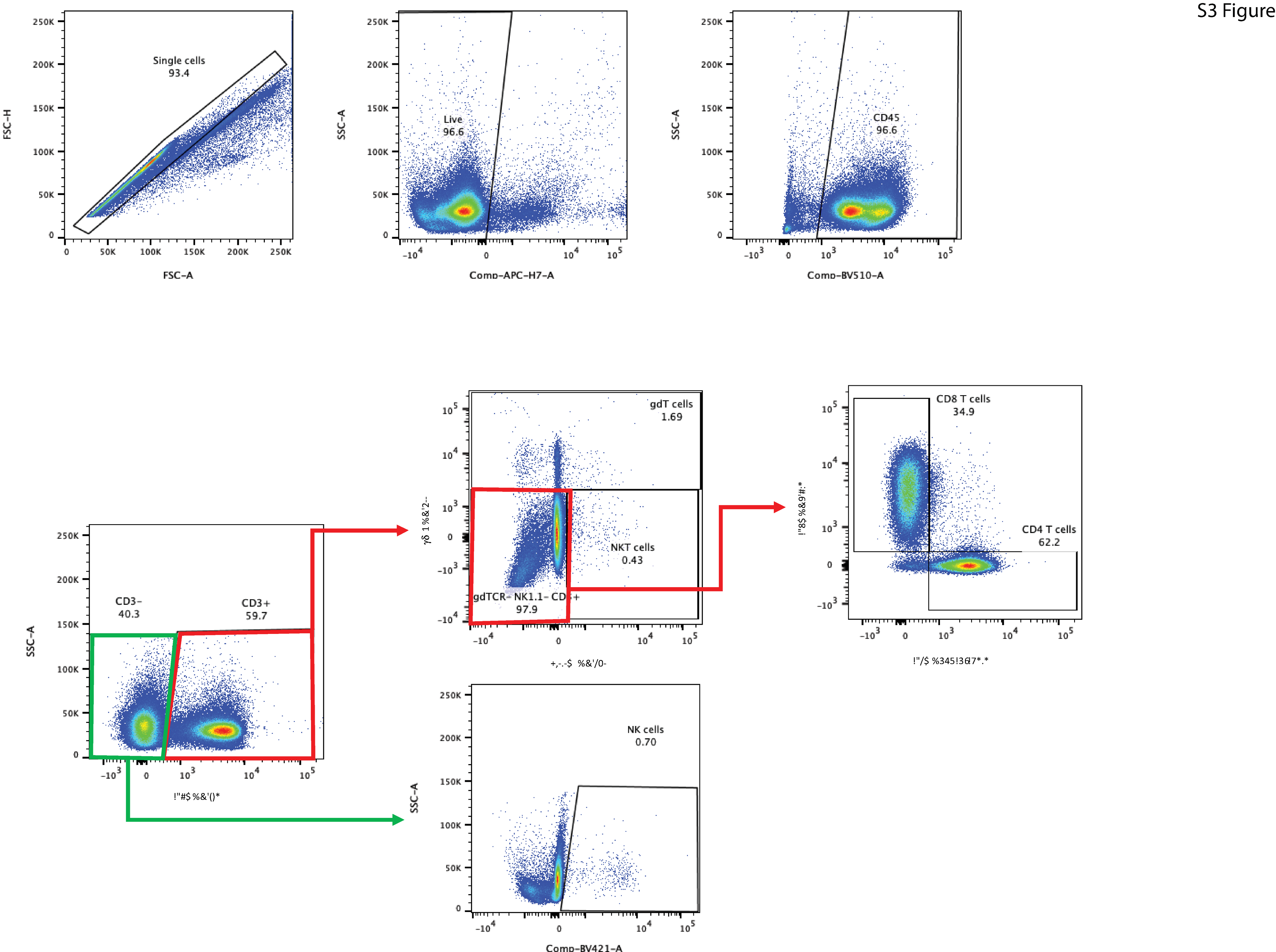

### Supplemental Figure 4

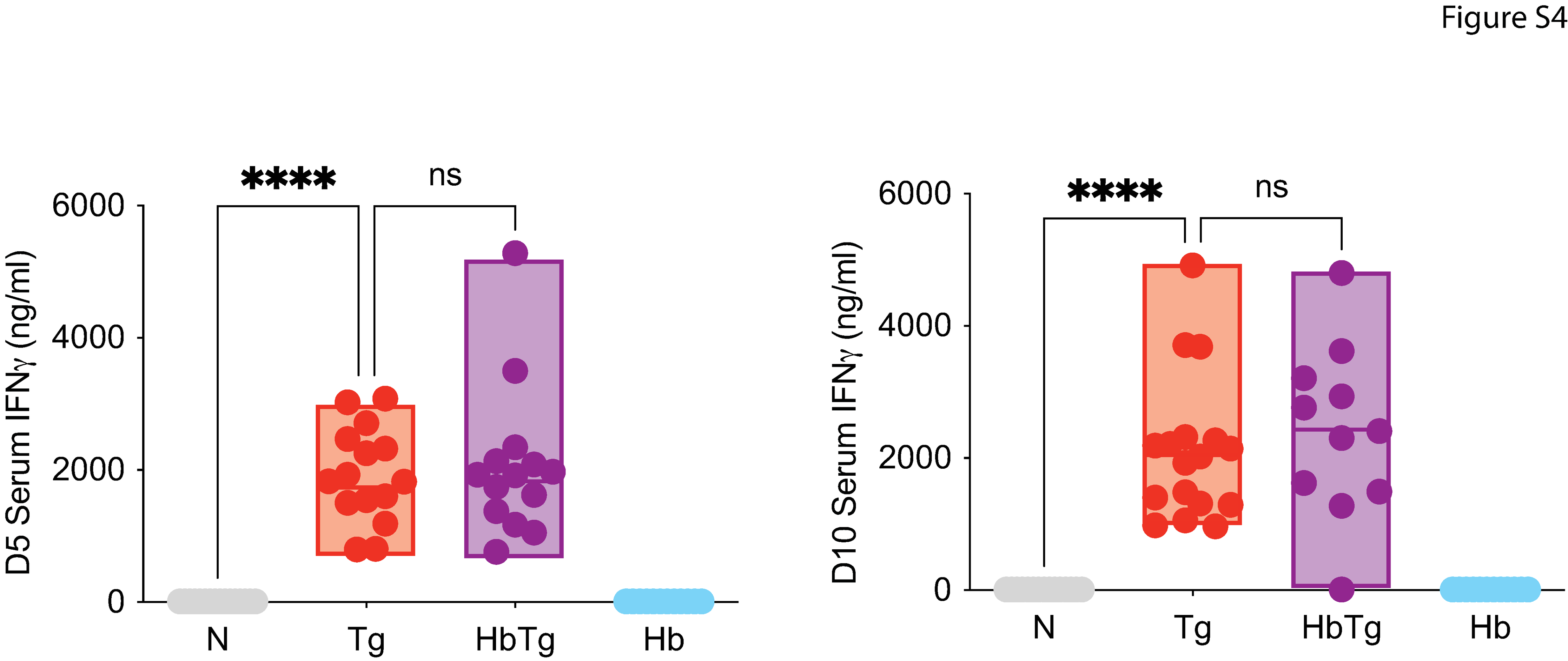

### Supplemental Figure 5

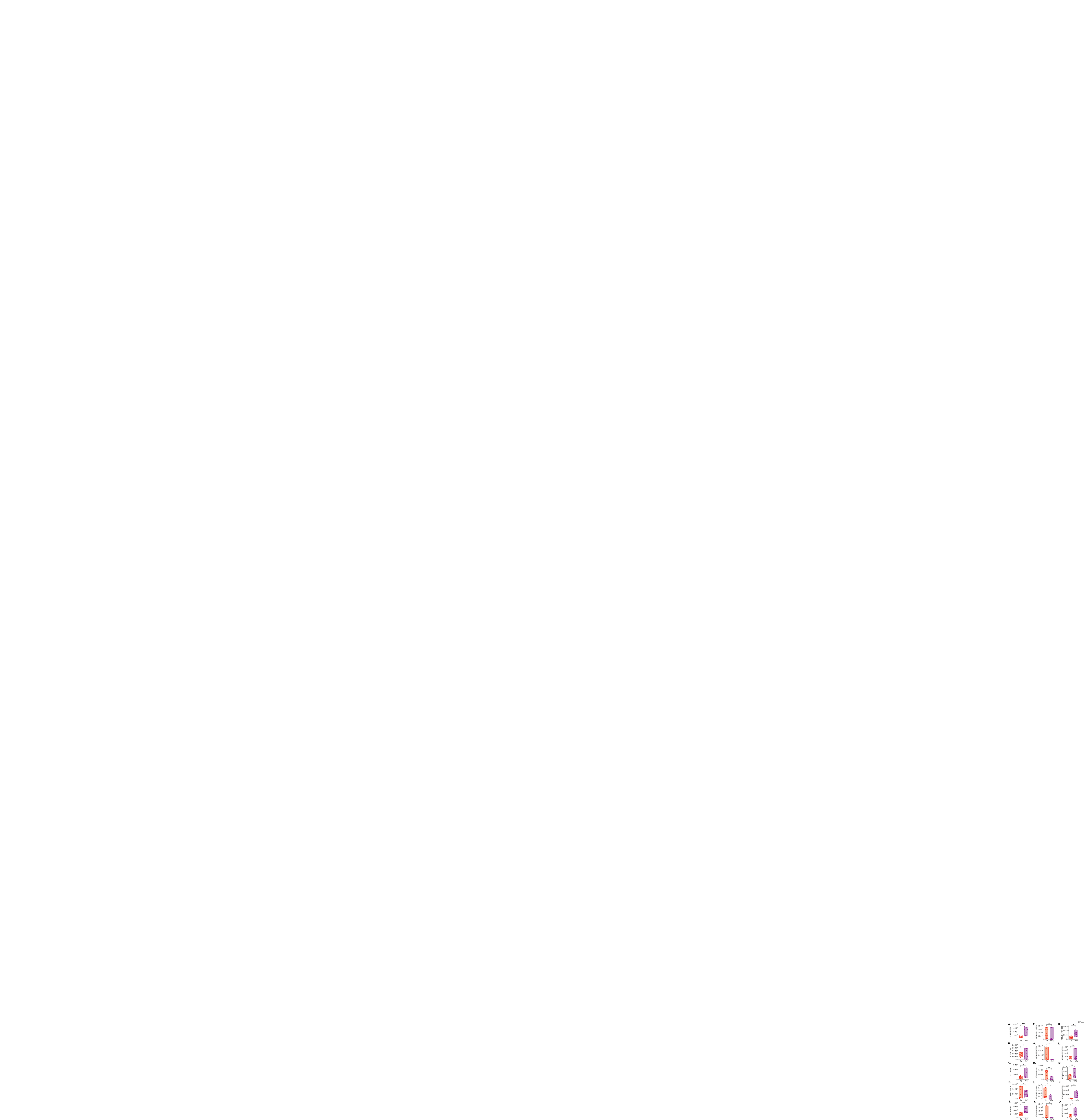

### Supplemental Figure 6

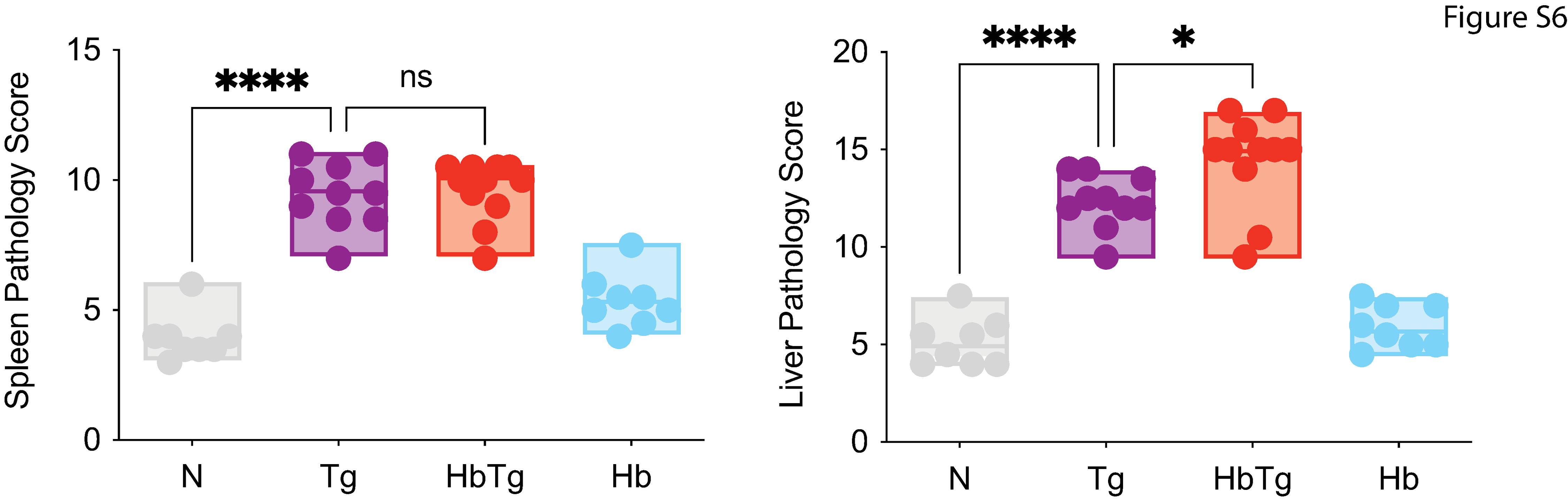
